# Insights into Molecular Diversity within the FET Family: Unraveling Phase Separation of the N-Terminal Low Complexity Domain from RNA-Binding Protein EWS

**DOI:** 10.1101/2023.10.27.564484

**Authors:** Courtney N. Johnson, Kandarp A. Sojitra, Erich J. Sohn, Alma K. Moreno-Romero, Antoine Baudin, Xiaoping Xu, Jeetain Mittal, David S. Libich

## Abstract

The FET family proteins, which includes FUS, EWS, and TAF15, are RNA chaperones instrumental in processes such as mRNA maturation, transcriptional regulation, and the DNA damage response. These proteins have clinical significance: chromosomal rearrangements in FET proteins are implicated in Ewing family tumors and related sarcomas. Furthermore, point mutations in FUS and TAF15 are associated with neurodegenerative conditions like amyotrophic lateral sclerosis and frontotemporal lobar dementia. The fusion protein EWS::FLI1, the causative mutation of Ewing sarcoma, arises from a genomic translocation that fuses the low-complexity domain (LCD) of EWS (EWS^LCD^) with the DNA binding domain of the ETS transcription factor FLI1. This fusion not only alters transcriptional programs but also hinders native EWS functions like splicing. However, the precise function of the intrinsically disordered EWS^LCD^ is still a topic of active investigation. Due to its flexible nature, EWS^LCD^ can form transient interactions with itself and other biomolecules, leading to the formation of biomolecular condensates through phase separation – a mechanism thought to be central to the oncogenicity of EWS::FLI1.

In our study, we used paramagnetic relaxation enhancement NMR, analytical ultracentrifugation, light microscopy, and all-atom molecular dynamics (MD) simulations to better understand the self-association and phase separation tendencies of EWS^LCD^. Our aim was to elucidate the molecular events that underpin EWS^LCD^-mediated biomolecular condensation. Our NMR data suggest tyrosine residues primarily drive the interactions vital for EWS^LCD^ phase separation. Moreover, a higher density and proximity of tyrosine residues amplify the likelihood of condensate formation. Atomistic MD simulations and hydrodynamic experiments revealed that the tyrosine-rich N and C-termini tend to populate compact conformations, establishing unique contact networks, that are connected by a predominantly extended, tyrosine-depleted, linker region. MD simulations provide critical input on the relationship between contacts formed within a single molecule (intramolecular) and inside the condensed phase (intermolecular), and changes in protein conformations upon condensation. These results offer deeper insights into the condensate-forming abilities of the FET proteins and highlights unique structural and functional nuances between EWS and its counterparts, FUS and TAF15.

## Introduction

The FET family of RNA binding proteins, Fused in Sarcoma (FUS), RNA binding protein EWS (EWS), and TATA-binding protein-associated factor 2N (TAF15), are predominantly nuclear proteins essential for transcription and mRNA processing^1–4^. Mutations within FET proteins are linked to pathogenesis in neurodegenerative diseases and cancers. Chromosomal translocations that fuse a FET N-terminal low-complexity domain (LCD) with a transcription factor-derived DNA binding domain (DBD) form potent oncogenic fusion proteins that are the driver mutations that cause Ewing sarcoma (EwS) and related malignancies^5–7^. In EwS, an aggressive soft tissue and bone cancer that is primarily diagnosed in pediatric patients, the self-associative and phase separation properties imparted by the EWS LCD (EWS^LCD^) in the archetypical FET fusion EWS::FLI1 are essential for neoplastic transformation and progression of the tumor^8–11^.

The phase separation properties of FUS, the exemplar member of the FET family, have been studied extensively while similar properties of EWS and TAF15 have been less thoroughly explored^12–15^. Consequently, much of our understanding about EWS phase separation has been inferred from studies of FUS. Previous work demonstrated phase separation of full-length FUS and the FUS LCD (FUS^LCD^) occurs due to multivalent interactions involving multiple residue types^16, 17^. Like the FUS^LCD^, the EWS^LCD^ is comprised of an abundance of serine, tyrosine, glycine, and glutamine residues that can establish multivalent interactions, leading to the formation of phase-separated droplets^13^ (**Fig. 1**). Further, modulation of FET LCD-LCD multivalent interactions can impact the regulation of transcription in fusion proteins^18^. Therefore, a detailed, molecular understanding of the self-associative and phase separation properties mediated by the EWS^LCD^ are essential for exploring the convergent and divergent roles of the FET proteins.

**Figure 1.**
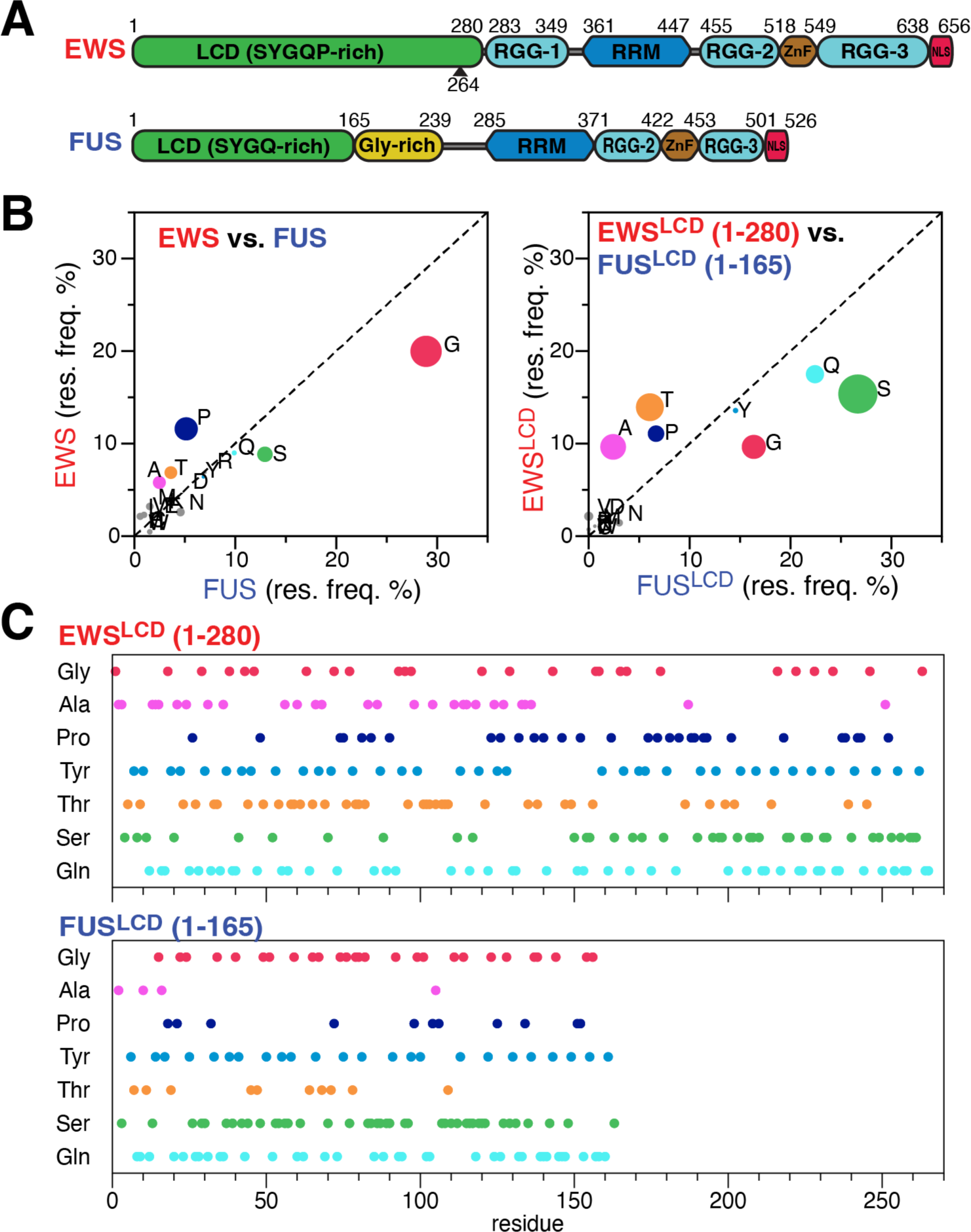
Comparison of (A) FUS and EWS domain architecture, numbers indicate domain boundaries, arrow marks the portion of EWS^LCD^ contributed to EWS::FLI1. (B) Percent abundance of amino acid in full-length (left) and the LCD (right) EWS and FUS, circle size denotes percent enrichment. (C) Amino acid distribution of the predominant amino acids in the EWS^LCD^ (top) and FUS^LCD^ (bottom).

Here we present a comprehensive, atomistic study of the phase separating properties of the EWS^LCD^ using paramagnetic relaxation enhancement (PRE) NMR, analytical ultracentrifugation, microscopy, and molecular dynamics simulations. Structural characterization of EWS^LCD^ reveal it is predominantly disordered at physiological conditions. Atomistic simulations and mutational analysis reveal tyrosine residues are the predominant force driving phase separation. Our experiments uncover ensemble conformational preferences for the N-terminal (less compact) and the C-terminal (more compact) regions of EWS^LCD^, findings that contrast the behavior observed in the FUS^LCD^. Importantly, through comparative analysis of biophysical experiments and molecular simulations of EWS^LCD^ in dilute and condensed phases, we find that interactions observed in the dilute phase inform on atomic interactions driving phase separation. The results of this study unveil the molecular framework governing phase separation of EWS^LCD^, expand our understanding of the phase separation of FET proteins, and provide insights useful for understanding the pathogenicity of oncogenic fusion proteins such as EWS::FLI1.

## Materials and Methods

### Protein Purification

Synthetic DNA sequences codon optimized for expression in *E. coli* coding for the N-terminal 264 residues of EWS (EWS^LCD^) and the tyrosine mutants EWS^LCD,7YS^ and EWS^LCD,13YS^ were obtained from GenScript (NJ). Single point mutations were introduced by site directed mutagenesis using the primers listed in **Supp. Table 1**. All constructs were cloned into a modified pET expression vector harboring an N-terminal His_8_-tag followed by a tobacco etch virus (TEV) cleavage site. The sequences of all constructs were confirmed by DNA sequencing (Genewiz). A list of all EWS^LCD^ mutants used in this study is provided in **Supp. Table 2**. Cleavage of the affinity tag with TEV leaves a two residue (Gly, Ala) cloning scar that are not included in the numbering scheme. Protein purification followed the procedure previously described^19^. Briefly, plasmids were transformed into *E. coli* BL21 DE3 Star^TM^ and grown in either Luria Broth (LB) or M9 media supplemented with 1 g/L ^15^NH_4_Cl and 0.02% (w/v) yeast extract at 37 °C. Recombinant protein expression was induced when the cultures reached OD_600_ ∼ 0.6 with 0.5 mM IPTG and continued for 3.5 hours at 37 °C. Cells were harvested by centrifugation, rapidly frozen, resuspended in 8 M urea, 20 mM CAPS pH 11, and lysed by passage through a high-pressure homogenizer (Avestin). The lysate was cleared by centrifugation at 45,000 g for 30 minutes, filtered (0.45 µm syringe filter), and loaded on a IMAC column. The protein was eluted with 300 mM imidazole, concentrated with 3 kDa cut-off spin concentrator (MilliporeSigma), diluted to 20 mL in 20 mM Tris pH 8.5, 1 mM 1,6-hexanediol and the His8 tag was cleaved with TEV at 20 °C overnight. Cleaved His_8_-tag and excess TEV were removed by passing the solution over a second IMAC column equilibrated in the same buffer, the flow through was concentrated to < 5 mL and applied to a HiLoad 16/600 Superdex 75 pg size exclusion column (SEC, Cytiva) equilibrated in 20 mM CAPS pH 11. Fractions containing the protein were pooled, concentrated to 1 mM in 20 mM CAPS pH 11, aliquoted and flash frozen. For constructs containing cysteines 1 mM TCEP was included in all buffer solutions.

### Fluorescent Labeling

Purified EWS^LCD^ was fluorescently labeled using sortase-mediated conjugation^20^. The N-hydroysuccinimide (NHS) activated DyLight 488 fluorescent dye (ThermoFisher) was solubilized in dimethylformamide and incubated with the SortC1 peptide, KLPETGG following the manufacturers protocol. Purified EWS^LCD^ was mixed with an equal concentration of fluorescently labeled SortC1 and 2.5 µM recombinant sortase^21^ in 100 mM sodium bicarbonate at pH 9.5, incubated overnight at room temperature, protected from light. The protein was isolated from free dye, peptide and enzyme by SEC using a HiLoad 16/600 Superdex 75 pg column. Fluorescently labeled protein, EWS^LCD,488^ was concentrated to 1 mM and frozen at −80 °C.

### Microscopy and Turbidity Assays

Stock EWS^LCD^ or mutants were diluted into low or high salt 20 mM Tris buffer pH 7.5 in a 96 flat-bottom microwell plate (Greiner Bio-One) and sealed with ultra-clear tape. Brightfield images and A_320_ measurements were simultaneously recorded with a BioTek Cytation Gen 5 imaging plate reader equipped with a 20x objective and phase contrast optics. For fluorescent images, 4 µL of EWS^LCD^ stock was mixed with 1% (v/v) fluorescently labeled EWS^LCD,488^, applied to a chambered coverslip (Grace Bio-Labs) coated with 1% Pluronic F-127, sealed with a second coated coverslip, and imaged with an Olympus FV300 inverted confocal microscope using a 40x oil immersion objective lens operating at 1% power using the 488 nm laser for transmitted light and fluorescence imaging. Fluorescence recovery after photobleaching (FRAP) was recorded on circular regions of interest irradiated at 10% laser power. Fluorescence intensity of bleached regions for which the contrast had been globally adjusted was obtained using FIJI^22^.

### Temperature Sensitivity

EWS^LCD^ samples were prepared in a 1.5 mL microcentrifuge tube at room temperature, transferred to a small volume (40 µL) quartz cuvette, sealed with parafilm, and the absorbance spectrum from 200-320 nm was measured using a diode array spectrophotometer outfitted with a Peltier unit for temperature control (Agilent). The measurements began at 25 °C, decreased to 10 °C then increased back to 40 °C. Room temperature N_2_ gas was passed over the outside of the cuvette to prevent condensation from forming on the cuvette during cooling steps.

### Pelleting and Soluble Fraction Assays

To measure the amount of protein that stays in solution due to increasing salt concentration, equal volumes of a 1 mM EWS^LCD^ in CAPS pH 11 were pipetted into aliquots of 20 mM Tris pH 7.5 varying (10 – 750 mM) concentrations of sodium chloride to reach a final protein concentration of 50 μM. To measure the amount of protein remaining in solution as a result of increasing EWS^LCD^ concentration, the volumes of 1 mM EWS^LCD^ were diluted in 20 μL CAPS pH 11 such that the total amount of 20 mM CAPS present in each 50 μL solution was the same regardless of the final protein concentration. Protein solution was mixed into 33.3 mM Tris buffer pH 7.4 so that the final concentration of Tris buffer was 20 mM. To measure the effect of mutating tyrosine residues on droplet formation, EWS^LCD^ and mutant samples were prepared in triplicate by diluting stock solutions into 20 mM Tris pH 7.5, 150 mM NaCl in 1.5 mL microcentrifuge tubes. Samples were agitated gently by hand to ensure proper mixing, promptly pelleted by centrifugation at 18,000 g for 15 min at room temperature, then carefully removed from centrifuge to prevent remixing. The absorbance at 280 nm of the supernatant was collected in triplicate for each sample on a Implen nanophotometer.

### Analytical Ultracentrifugation

Single wavelength (280 nm) sedimentation velocity experiments were recorded with 450 µL samples in 2-sector epon centerpieces with quartz windows on an Optima AUC instrument at 20 °C and 35,000 rpm. Data were collected in intensity mode and analyzed with Ultrascan (version 6731) Two-dimensional spectrum analysis was performed following standard protocols (Demeler, 2010). The van Holde-Weishet analysis function in Ultrascan was used to generate plots of G(s) integral distributions of the sedimenting species vs normalized concentrations of the boundary signal.

### Size Exclusion Chromatography

Experiments were carried on an analytical Superdex 200 10/300 size exclusion chromatography column (Cytiva) equilibrated with either 20 mM CAPS, pH 11.0 or 20 mM MES pH 6.0. Samples of 200 µL of increasing concentrations of EWS^LCD^ (10 µM, 20 µM, 40 µM, 100 µM, 200 µM) were injected and eluted at a flowrate of 0.4 mL/min. Absorbance was measured at 280 nm.

### Nuclear Magnetic Resonance

NMR experiments were conducted on a Bruker Avance NEO spectrometer operating at a proton Larmor frequency of 700.13 MHz at 25°C. Data were processed using NMRPipe^23^ and analyzed with Sparky^24^. ^1^H,^15^N-HSQC, *T*_1,_ *T*_1ρ,_ and heteronuclear NOE experiments were recorded on 50 µM EWS^LCD^ in 20 mM MES pH 5.5. ^15^N *R*_1_ and *R*_2_ relaxation rates were calculated from *T*_1_ and *T*_1ρ_ experiments using 256 x 1024 complex data points in the indirect (^15^N) and direct (^1^H) dimensions which correspond to 81.9 and 112.6 ms. The ^15^N *T*_1_ experiment consisted of 8 interleaved spectra with relaxation delays of: 40, 80, 120, 200, 280, 400, 600, 800 ms. The *T*_1ρ_ experiment was recorded using a *B*_1_ field of 1400 Hz and 8 interleaved spectra with relaxation delays of: 1, 21, 31, 41, 61, 81, 121, 201 ms. ^15^N *R*_2_ rates were calculated using equation 1:

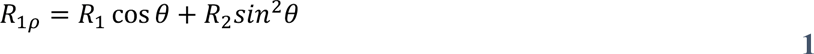

where θ = arctan(ω_1_/Ω), ω_1_ is the *B*_1_ field strength and Ω is the offset from the spinlock carrier frequency. ^1^H-^15^N heteronuclear NOE experiments were recorded on the same samples as those used for relaxation experiments. The ^1^H-^15^N heteronuclear NOE experiment consists of two interleaved experiments, plus and minus proton saturation, each recorded using a recycle delay of 4 seconds. Spectra were acquired with 64×1024 complex points in the indirect (^15^N) and direct (^1^H) dimensions which correspond to 41.0 and 112.6 ms.

### Diffusion Spectroscopy

Experiments were recorded on a 50 µM sample of EWS^LCD^ dissolved in either 20 mM CAPS, pH 11.0 or 20 mM MES pH 6.0. The diffusion time 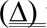 was set to 0.1 s and the gradient pulse length 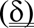 to 3 ms for 20 different gradient strengths: 0.96, 3.39, 5.83, 8.26, 10.69, 13.12, 15.55, 17.98, 20.42, 22.85, 25.28, 27.71, 30.14, 32.58, 35.01, 37.44, 39.87, 42.3, 44.74, and 47.17 G/cm. Intensity ratios were calculated by fitting of the ^1^H spectrum region corresponding to the aromatic side-chains (between 7.3 and 6.6 ppm).

### Paramagnetic Relaxation Enhancement

The nitroxide spin label, 3-Maleimido-2,2,5,5-tetramethyl-1-pyrrolidinyloxy (5-MSL), was conjugated to the EWS^LCD^ via maleimide chemistry at non-native cysteines (S40C, S116C, S149C, S239C, or S260C). A selected purified EWS^LCD^ cysteine mutant was exchanged into degassed buffer containing 2 M urea, 50 mM Tris pH 7 and 1 mM TCEP and incubated with 5-MSL (50 mg/mL in 95% ethanol) at a 20x molar ratio for 60 hours at room temperature with gently nutation while protected from light. To remove excess non-reacted spin label, the solution was applied to a Superdex 75 16/60 HiLoad size exclusion column equilibrated in 20 mM CAPS pH 11, fractions containing the spin-labeled EWS^LCD^ were concentrated to 1 mM in 20 mM CAPS pH 11, aliquoted and flash frozen for storage at −80°C. Samples were prepared by diluting an aliquot of 1 mM ^15^N and spin labeled EWS^LCD^ to a final concentration of 50 µM in 20 mM MES pH 5.5, 1 mM TCEP, 5% D_2_O in a 5 mm glass NMR tube. For recording solvent PREs, 2.75 μL of 100 μM 5-MSL was added directly into a 5 mm glass NMR tube containing 50 µM EWS^LCD^ (no native cysteines) for a final concentration of 0.5 μM 5-MSL representing 1% of the total protein concentration ^1^H_N_-*T*_2_ experiments were recorded using the two time-delay point approach^25^, (0.001 and 0.6 ms) with 120 x 1024 complex data points in the indirect (^15^N) and direct (^1^H) dimensions corresponding to 38.4 and 112.6 ms. The paramagnetic samples were subsequently quenched with a 10x molar excess of ascorbic acid and diamagnetic data were recorded using identical experimental parameters. The data was processed using NMRPipe^23^ and exported to Sparky^24^ (build 04/22/2021) for peak picking and extraction of peak intensities. The proton relaxation rate was calculated using equation 2:

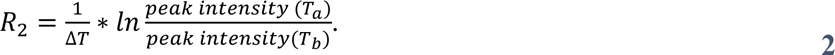

where ΔT = T_b_-T_a,_ here 0.6 ms. The difference between the paramagnetic relaxation rate and the diamagnetic relaxation rate is used to calculate the ^1^H_N_-*Γ*_2_ rates.

### Coarse-Grained (CG) Simulation

Single-chain CG simulation was performed using LAMMPS simulation package^26^ for 2 μs using the HPS-Urry model^27^. For CG co-existence simulations, HOOMD-blue 2.9.3 simulation software package^28^ was used following the protocol established in previous work^29,30^. The initial configuration was prepared with 100 chains of EWS LC using HPS Urry model in a slab of 17.5 nm x 17.5 nm x 122.5 nm. The simulations were performed at 300 K with timestep of 10 fs. Langevin thermostat was used with residue friction coefficient, γ = m_i_/t_damp_, where m_i_ is mass of each residue and t_damp_ is damping factor, which was set to 1000 ps. The mutant studies were performed at 320 K, since we observe co-existing phases for all the systems at this temperature, which allows us to compare their saturation concentrations. Each simulation was run for 5 μs, and the analysis was conducted by excluding the initial 1 μs as equilibration time. Error bars were computed by dividing the trajectory into 4 blocks and estimating the standard error of mean between the blocks.

### All-Atom (AA) Simulation

AA Single-Chain: To obtain the initial atomistic structure of single-chain EWS^LCD^, CG simulations were performed. Based on the radius of gyration (*R*_g_) distribution, four different structures were selected near the mean *R*_g_. We then used Modeller^31^ to convert C_*α*_ coordinates from the CG simulation to AA coordinates. The system was then prepared using GROMACS 2020^32^ using an optimized protein force field, Amber99SBws-STQ, with improved residue-specific dihedral correction^33^ and TIP4P/2005 water model^34^. At first, the single chain EWS^LCD^ was placed in an octahedral box of 13.5 nm size. Vacuum minimization was performed followed by solvating the protein using TIP4P/2005 water molecules. The system’s energy was further minimized after solvation to relax any overlaps between the protein and water atoms. To maintain a neutral charge and achieve a salt concentration of 100 mM, sodium ions (Na+) and chloride ions (Cl-) were introduced into the system. The improved salt parameters, as proposed by Luo and Roux^35^, were utilized for this purpose. After preparing the system, the temperature was equilibrated at 300 K in NVT (canonical ensemble) using Nose-Hoover thermostat^36^ with a coupling constant of 1.0 ps. To reach an equilibrium pressure of 1 bar, NPT (isothermal-isobaric) equilibration was carried out using Berendsen barostat^37^ with isotropic coupling with a coupling constant of 5.0 ps. The production runs were performed in NVT ensemble. The temperature was maintained at 300 K using Langevin middle integrator^38^ and 1 ps^−1^ friction coefficient was used. Four independent simulations starting with different initial configurations were performed using OpenMM7.5^39^ for 3.75 *μ*s each. We use the Hydrogen mass repartitioning scheme by setting its mass to 1.5 amu to use a larger 4 fs timestep^40^. Long-range electrostatic interactions were calculated using the Particle Mesh Ewald (PME) method^41^. For short-range van der Waal interactions, the cutoff was set to 0.9 nm. SHAKE algorithm^42^ was used to apply constraints to all hydrogen containing bonds.

The *R*_g_ autocorrelation function (ACF) was evaluated for each system to calculate the correlation time where the ACF decays to 1/e. The average correlation time for the four systems was ∼ 85 ns. Based on the correlation time, we removed the first 100 ns as the time the system took to equilibrate from the arbitrary initial condition and carried out the analysis for the remaining simulation time.

For tyrosine mutant systems, 7YS and Y170S/Y172S, total simulations time over four replicas were 8 *μ*s and 6 *μ*s respectively. To compute contact maps, we considered two heavy atoms as forming a nonbonded contact if the distance between them was less than 0.6 nm. The total contact between two residues, i and j, was determined as the cumulative sum of all contacts formed between any heavy atom in the i^th^ residue and any heavy atom in the j^th^ residue. This approach effectively accounted for the contribution from sidechains, with longer sidechains having a greater potential for interaction. To assess the simulation results against experimental PRE data, we determined the <r^−6^>/<r^−6^>_max_, where r represents the distance between two heavy atoms within a 3.5 nm cutoff distance. The error bars were computed by estimating the standard error of mean over all the replicas of a particular system. The trajectories were examined and the TTClust python program was used to extract ten clusters^43^.

### AA Condensed Phase Simulation

For preparing AA slab simulation, we use a similar method as described previously^44^. Briefly, to obtain the initial structure for multichain simulation, 25 chains of the EWS^LCD^ were first equilibrated at 300 K using CG co-existence simulation. The AA system was made from the C_**α**_ positions of the CG slab using Modeller. Clashes between the side chains were removed with CAMPARI Monte Carlo engine and ABSINTH implicit solvent model with a fixed backbone^45^. Using this conformation, soft core minimization was performed followed by steepest descent energy minimization in GROMACS 2020. After minimization, the system was solvated, ionized, and equilibrated, following similar steps used for single chain AA simulation. The production run was performed for 1 µs and 200 ns for equilibration were removed prior to analysis.

### Calculation of NMR Relaxation Parameters

To analyze the NMR relaxation parameters we employed a similar method to that described in a prior study^46^. The trajectories were divided into blocks of 100 ns to compute N-H bond vector autocorrelation for each residue of each trajectory. The average autocorrelation decay over all the 100 ns blocks is then fitted to a second-order exponential function (eq. 3). Using the fitting parameters, the spectral density function (eq. 4) is computed, which is then used to compute relaxation rates.

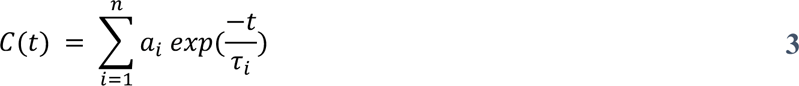

where n = 2 for our case.

We obtain spectral density function by computing the analytical Fourier transform of the autocorrelation function:

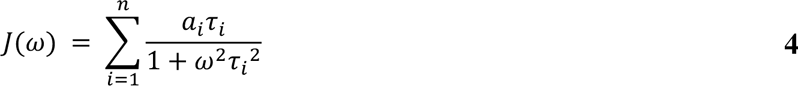

We obtain *R*_1_ (eq. 5), *R*_2_ (eq. 6) and NOE (eq. 7) as a function of spectral density function sampled at eigen frequencies of the spin system:

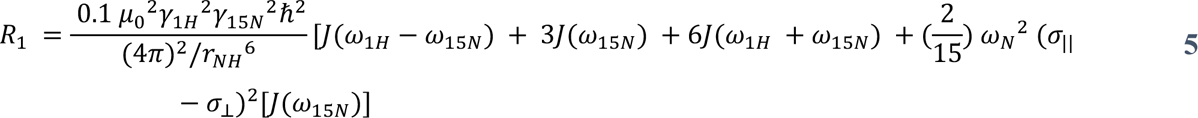

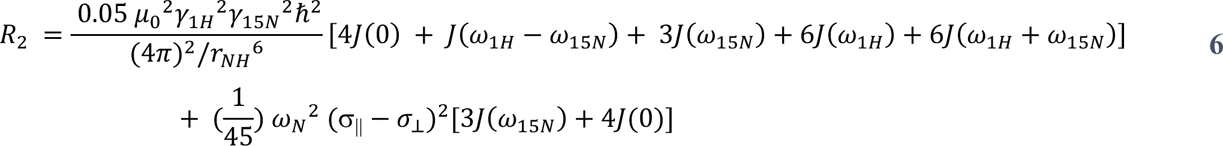

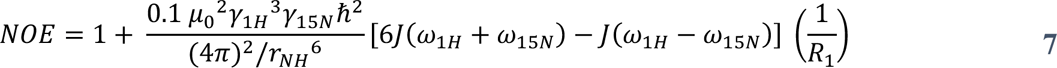

where μ_0_ is the permeability of free space, γ_i_ is gyromagnetic ratio of the spin i, *ℏ* is reduced Planck’s constant, α_||_ and α_⊥_ re parallel and perpendicular components of the axially symmetric ^15^N chemical shift tensor, respectively. Value of α_||_ - α⊥ is taken as −163 ppm^47^, internuclear ^1^H – ^15^N distance, r_NH_ is 1.02 *Å* and 700.15 MHz ^1^H Larmour frequency is assumed for the calculation.

## Results

### EWS Domain Architecture and Differences between FUS and EWS

The FET family of RNA binding proteins is defined by conserved domain architecture that includes an amino terminal low-complexity domain (LCD), RGG repeats, RNA recognition motif, a zinc finger, and a carboxy terminal nuclear localization signal (**Fig. 1A**). The similarities of FUS and TAF15 extend beyond the domain architecture as they share very similar amino acid composition in the structured domains as well as in the N-terminal LCD (**Supp. Fig. 1A**). In contrast, EWS has distinct differences in both amino acid composition and distribution when compared with FUS and TAF15. Full-length EWS is enriched in alanine, threonine, and proline and depleted in glycine when compared to FUS and TAF15 (**Fig. 1B, Supp. Fig. 1A**). The difference in amino acid composition is more acute when comparing the LCDs, FUS^LCD^ and TAF15^LCD^ are enriched in glycine, serine, and glutamine relative to EWS^LCD^ (**Fig. 1B, Supp. Fig. 1B**). Further stratifying the differences between EWS^LCD^ and FUS^LCD^, the distribution of alanine and serine are predominantly clustered in the N- and C-terminus of EWS^LCD^ respectively while in FUS^LCD^ serine is evenly distributed across the sequence (**Fig. 1C**). Moreover, the tyrosine appears to be relatively well distributed in FUS^LCD^, while in EWS^LCD^ there is a patch between residues 128 to 157 that is devoid of tyrosine (**Fig. 1C**). These observations clearly highlight the dissimilarities in amino acid distribution and composition between FUS and EWS. Despite these disparities, FUS has remained a model protein in understanding LLPS and the formation of biomolecular condensates of the FET family^15, 17, 48-50^.

### Physiochemical Properties of EWS^LCD^

Turbidity measurements reveal that EWS^LCD^ droplet formation occurs under a pH range from 5 to 9 suggesting that charge-dependent interactions may not be the primary driving force for self-association (**Fig. 2A**). However, turbidity was observed to increase linearly with increasing NaCl concentrations and were found to positively depend on the concentration of the EWS^LCD^ (**Fig. 2B**), which is consistent with the “salting-out” behavior shown by other LCDs^51, 52^. Sample turbidity increases as the temperature is decreased, and upon heating the sample becomes clear (**Fig. 2C**) highlighting upper critical solution temperature (UCST) behavior of EWS^LCD 53^. Microscopy using fluorescently tagged EWS^LCD,488^ revealed the presence of protein droplets that spontaneously condense and fuse to form large droplets based on the solution concentration, an effect that is enhanced by the presence of 0.15 M NaCl (**Fig. 2D**). EWS and EWS^LCD^ have previously been reported to undergo LLPS forming biomolecular condensates^1, 12, 13, 54^, and the results presented here are consistent with these reports. Pelleting assays conducted on increasing concentrations of EWS^LCD^ in 20 mM Tris pH. 7.4, 150 mM NaCl reveal that under these conditions, at 25 µM total EWS^LCD^, more than two-thirds is being incorporated into condensates, a trend that continues to the highest measured EWS^LCD^ concentration of 400 µM (**Fig. 2E**). Similarly, in pelleting assays 50 μM EWS^LCD^ is soluble when NaCl is excluded from the solution but begins to form condensates with the addition of 100 mM NaCl and with increasing NaCl concentration, more EWS^LCD^ is driven into the condensed phase (**Fig. 2F**). Co-existence CG simulations of EWS^LCD^ in slab geometry also highlight the formation of co-existence phases (**Supp. Fig. 2A**). Moreover, the simulations support the experimental observations, that compared to FUS^LCD^, EWS^LCD^ forms droplets at lower protein concentrations and its critical temperature (temperature above which a single phase is observed without droplet formation) is higher (**Fig. 2G, H**).

**Figure 2.**
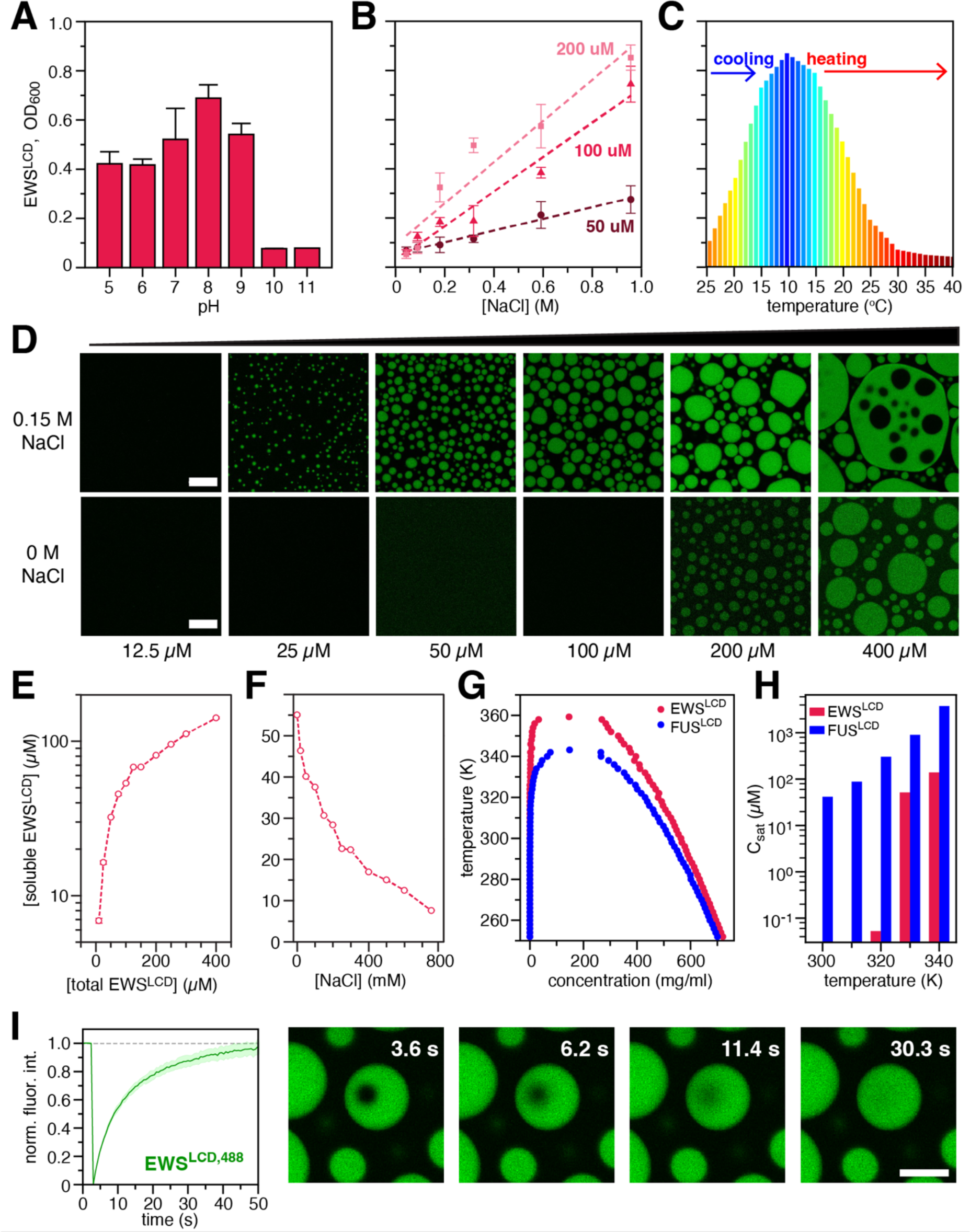
Phase separation propensity of EWS^LCD^ at varying (**A**) pH; (**B**) NaCl concentration; and (**C**) temperature. (**D**) Microscopy of EWS^LCD^ condensates in 20 mM Tris pH 7.4, with 150 mM (top panels) or without NaCl (bottom panels). Pelleting assays varying (**E**) EWS^LCD^, or (**F**) NaCl concentration. (**G**) Phase diagram and (**H**) *C*_sat_ comparison for FUS^LCD^ and EWS^LCD^ calculated from co-existence CG simulations. (**I**) FRAP of EWS^LCD^ condensates in 150 mM NaCl. Unless noted otherwise, all samples were 50 μM EWS^LCD^ in 20 mM Tris, pH 7.4 at room temperature. Scale bars are 10 µm.

Multiple biophysical methods were used to examine the degree of extension of EWS^LCD^ in 20 mM CAPS pH 11 and 20 mM MES pH 5.5. Size exclusion chromatography experiments show a lower retention volume of EWS^LCD^ in CAPS pH 11 compared to in MES pH 5.5, suggesting a higher compaction of the protein in the latter buffer (**Supp. Fig. 2B, 2C**). Sedimentation velocity analytical ultracentrifugation revealed that the sedimentation coefficient of EWS^LCD^ is larger in 20 mM MES pH 5.5 than in 20 mM CAPS pH 11, due to the decreased friction on the molecule as it begins to collapse onto itself through self-association in neutral buffers (**Supp. Fig. 2D, 2E**). Finally, diffusion ordered spectroscopy (DOSY) experiments show a faster translational diffusion of EWS^LCD^ in 20 mM MES than in 20 mM CAPS pH 11 (**Supp. Fig. 2F**). Taken together, these results show that EWS^LCD^ is more extended in 20 mM CAPS pH 11 as in 20 mM MES pH 5.5.

The liquid-like nature of EWS^LCD^ droplets was explored using fluorescence recovery after photobleaching (FRAP) that showed the fluorescence intensity of the bleached spot rapidly recovered to almost 100% of the initial intensity in less than one minute indicative of a liquid state (**Fig. 2I**) ^55^. After aging the droplets for 24-72 hours the fluorescence intensity of the bleached spots does not recover completely as compared to the freshly formed droplets indicating that they have changed to become less liquid-like, a phenomenon referred to as “aging” and consistent with the reported behavior of other systems^56^ (**Supp. Fig. 2G**). Compared with the well-established physicochemical behavior of FUS^LCD^, we find that EWS^LCD^ is more prone to phase separate (lower *C*_sat_), which is expected from its higher multivalency (longer chain length) with a similar average composition of aromatic tyrosine residues (**Fig. 1B**).

### Peptide Chain Dynamics and Self-Association of the EWS^LCD^

We use NMR and molecular simulations to provide insights into the dynamics and identify the interactions of the EWS^LCD^ residues that are important for mediating phase separation. In comparison with FUS and other IDPs that have been studied in the condensed phase^17, 57^, here, NMR experiments were recorded in conditions under which the EWS^LCD^ was in the dilute phase. Our attempts to reconstitute condensed phase EWS^LCD^ NMR samples resulted in samples that rapidly aggregated when contrasted with our successful reconstitution of condensed phase FUS^LCD^ following previously reported protocols^17^. This further highlights differences in the two proteins that may have otherwise been considered similar in all regards.

Consistent with secondary structure predictions derived from chemical shift data^19^, the ps-ns dynamics reported by the *R*_1_, *R*_2_ and hetNOE experiments do not reveal any regions of stabilized structure and reveal a dynamic and disordered polypeptide. Specifically, the longitudinal relaxation rate (*R*_1_) profile is uniform with all residues having a similar value of ∼1.5 s^−1^ (**Supp. Fig. 3**). The ^1^H-^15^N hetNOE also has a flat profile with an average value of 0.25 s^−1^ (**Supp. Fig. 3**). Similarly, the transverse relaxation rate (*R*_2_) is predominantly uniform, consistent with the lack of secondary structure, however, an increase of ∼1 s^−1^ is observed for residues 160-210 and a similar decrease (∼1 s^−1^) is observed for residues 130-155 that are not reflected in the *R*_1_ and hetNOE profiles (**Supp. Fig. 3**). Relaxation rates from single chain atomistic simulations with 15 *μ*s total run time (**see Methods**) reproduces the salient features of the experimental data, which gives us confidence for using the simulated ensemble in interpreting the underlying molecular details. We note that the simulation rates with current state-of-the-art models are slower than experiment, which is consistent with other studies proposing reweighting strategies for obtaining a quantitative match between the two^58, 59^.

Paramagnetic relaxation enhancement NMR is sensitive to intra- and intermolecular interactions up to 30 Å away even if these interactions are transient and lowly-populated. Here we measured PREs to gain insight into the transient and long-range interactions that occur intra- and inter-molecularly that are important for mediating phase separation. We hypothesized that despite the dilute phase conditions under which the NMR was conducted, interactions important for phase separation would still occur, albeit at a much lower frequency^17, 48, 60^. To facilitate the conjugation of the paramagnetic spin label to the EWS^LCD^, which contains no native cysteines, serines S40, S116, S149, S239, and S260 were chosen as conservative mutation sites for the introduction of cysteine residues. With the 5-MSL spin label at S40C in the amino terminus intramolecular PREs of ∼ 10-15 s^−1^ are observed for residues 50-100, 110-130 and 170-190. Notably, lower PREs are observed in regions that are depleted of tyrosine residues, particularly 100-110 and 130-160 (**Fig. 3A**, dotted lines denote position of tyrosines). When the spin label is attached at S116C or S149C the overall magnitude of intramolecular PREs observed is uniformly ∼5-8 s^−1^without well defined stretches of higher rates except for in the immediately adjacent residues 170-185 (**Fig. 3A**). The large errors in these residues are due to the absolute broadening of these residues in the paramagnetic state. There is not an overall decrease in the peak intensity for all residues indicating there is no change in concentration due to precipitation. Additionally, there are no new peaks in the spectra that would be indicative of degradation. With the spin label in the C-terminal region at S239C, the large magnitude PREs (10-12 s^−1^) were only observed in the adjacent 160-210 and the region immediately adjacent to the PRE tag. When the spin label is placed at S260C larger PREs (10-12 s^−1^) were observed, similarly to the S239C mutant, in the regions 160-210 and 235-245. Enhancements in the range of 5-8 s^−1^ were observed across the entire chain and were correlated with the same residues as when the tag was at S40C (**Fig. 3A**).

**Figure 3.**
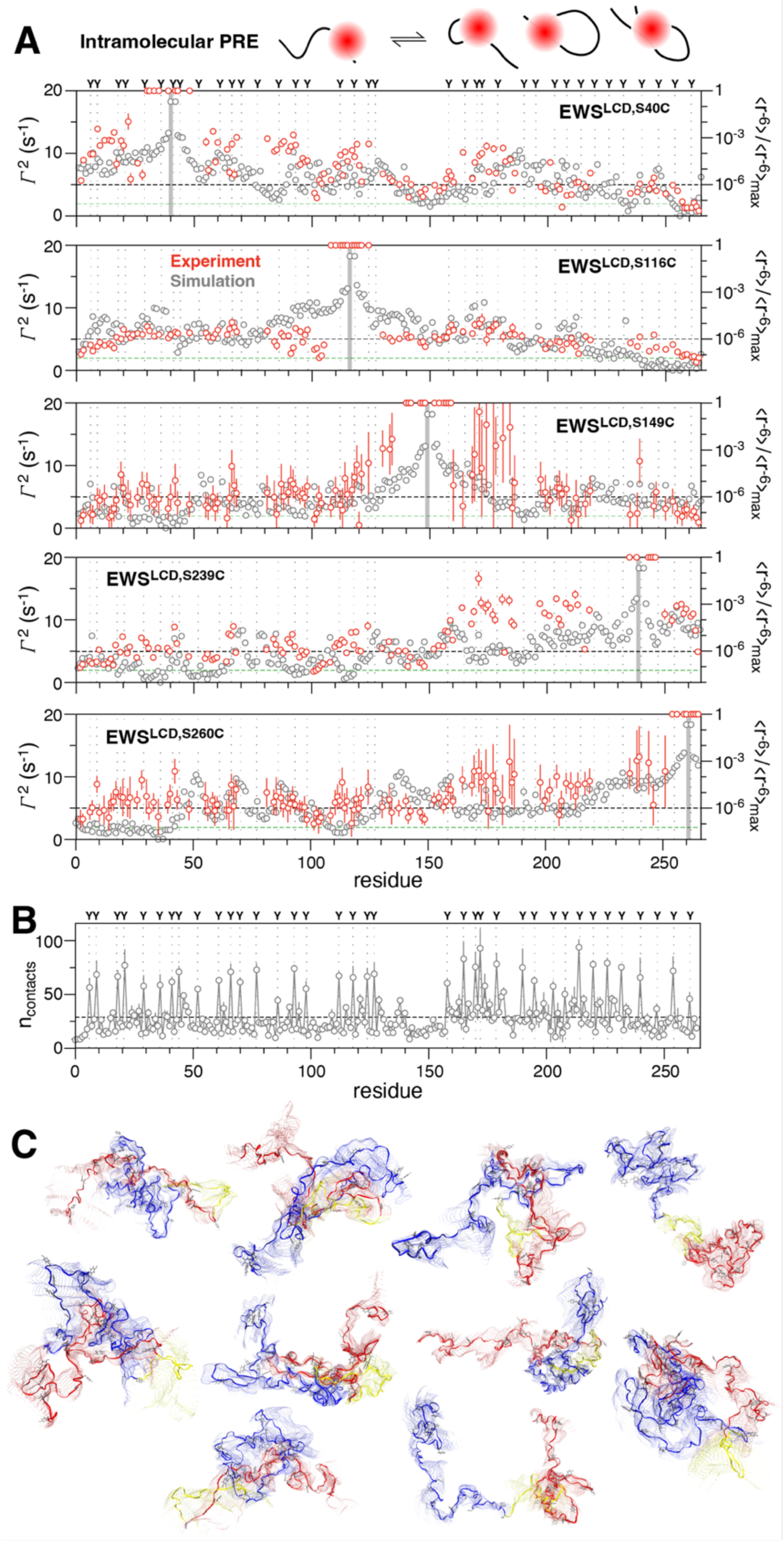
Intramolecular PRE rates (cartoon) measured for (A) EWS^LCD^ (red, left axis) with spin label placement (grey bar), position of tyrosine residues (dotted lines), and average solvent PRE (green dashed line) indicated. Black dashed line is drawn at 5 s^−1^ to guide comparison between panels. Intrachain contacts expressed as <r^−6^>/< r^−6^>_max_ (grey, right axis), (B) average per-residue contacts, and (C) selection of highly populated EWS^LCD^ ensemble images (N-terminus, 1-128, blue; middle linker 129-158, yellow; C-terminus, 159-265, red) extracted from single chain atomistic simulations.

Solvent PREs recorded with a soluble version of the 5-MSL tag show a flat profile of ∼2 s^−1^ across the entire length of the EWS^LCD^ indicating that the effects observed are due to protein-protein interaction and not arising from non-specific interactions of the protein with the solvent (**Supp. Fig. 4A**). Further, intermolecular PREs measured with natural abundance (e.g. predominantly ^14^N) 5-MSL labelled EWS^LCD^ S40C, S149C, and S260C mutants and unmodified ^15^N enriched EWS^LCD^ did not produce significant PREs beyond the ∼2 s^−1^ background (**Supp. Fig. 4B**). These data establish that the observed PREs arise from specific intramolecular contributions.

PRE values in molecular simulations can be determined through the DEER-PREdict software, which employs a rotamer library approach to position the paramagnetic probe’s conformation at the spin-labeled site^61^. By default, the MTSL 175 K rotamer library is utilized. While it is possible to incorporate new rotamer libraries, obtaining rotamer information for 5-MSL proved challenging. To bridge this gap, we calculated <r^−6^>/<r^−6^>_max_ from single-chain atomistic simulations. The resulting data exhibited a qualitatively similar trend to the PRE values (**Fig. 3A**). This correspondence provided us with the confidence to delve deeper into exploring intrachain interactions using single-chain atomistic simulations. The intrachain interactions suggests the C-terminal region forms the highest total number of contacts, and the linker region, devoid of tyrosine (130-160), forms the least (**Fig. 3B, Supp. Fig. 4C**). Interestingly, the total contacts show a similar trend as *R*_2_, which strongly suggests that the difference in contacts formed by the linker and C-terminal regions is causing their slower and faster relaxation, respectively (**Supp. Fig. 3A**). Simulation snapshots of some of the most populated structures (**Fig. 3C**), reveal the C-terminal and the N-terminal regions adopt compact conformations, while the linker region stays extended in most of the snapshots. To quantify the collapse of these regions, we computed the intrachain distance and *R*_g_ distribution for three segments of equal length (41-100, 101-160, 161-220). We observed that segment at C-terminal region (41-100) is more collapsed, followed by N-terminal region (101-160) and the linker region (161-220) (**Supp.** Figs. 4D,E).

To gain insights into the specific residue types contributing to intramolecular interactions, we analyzed pairwise residue type contacts. Our analysis considered all contacts formed between two heavy atoms of a residue type pair (see methods), providing a comprehensive assessment of the contribution of different residue types based on their interaction strength. Notably, our findings identified Tyr-Tyr interactions as the strongest. Furthermore, we observed that, in addition to Tyr-Tyr interactions, residue type pairs such as Tyr-Gln, Tyr-Ser, and Tyr-Pro also formed significant contacts (**Supp. Fig. 4F**). Consistent with prior research, our results underscore the role of transient interactions, including π-π, *sp*^2^/π, and hydrogen bonding involving various residue types, in stabilizing interactions within the dilute phase^17, 62^. In summary, our combined findings from NMR, PRE analysis, and molecular simulations provide a comprehensive perspective on EWS^LCD^, highlighting the influence of transient interactions in shaping intramolecular contacts.

### Tyrosine Residues Mediate Phase Separation of the EWS^LCD^

In vivo investigations have determined that the EWS^LCD^ is absolutely required for neoplastic transformation of Ewing sarcoma cells^8, 9, 63^. These studies identified that tyrosine depletion within the EWS^LCD^ through either mutant substitution (e.g. Y for S) or by truncation eliminated the oncogenic effect of EWS::FLI1 pointing to the importance of the self-associative, and possibly phase separation properties, of the EWS^LCD^ for oncogenesis. Indeed, our dilute phase experiments and simulations revealed the importance of tyrosine for self-association of EWS^LCD^ thus, the role of tyrosines within the EWS^LCD^ was investigated. Two tyrosine depleted mutants (EWS^LCD,Y170S/Y172S^ and EWS^LCD,7YS^) of the EWS^LCD^ were constructed along with the point mutants Y44S, Y208S, and Y172S (**Table 1**). Phase diagrams were constructed for EWS^LCD^ (**Fig. 4A**), EWS^LCD,Y170S/Y172S^, (**Fig. 4B**), and EWS^LCD,7YS^, (**Fig. 4C**) based on turbidity measurements. To investigate if distribution of aromatic amino acids has an outsized effect on their role in phase separation, Y170/Y172S was designed as it is the single YxY motif in EWS^LCD^. There are 4 YxxY motifs and all other tyrosines are separated by 3 or more residues. The 7YS mutant was designed to target YxxY and YxY motifs (**Supp. Table 1**). Phase diagrams constructed from turbidity measurements at OD_600_ show that phase separation of EWS^LCD^ begins at 25 µM protein and 100 mM NaCl (**Fig. 4A**). As the number of tyrosines is decreased the phase boundary shifts to the right; EWS^LCD,Y170S/Y172S^ does not form phase separated droplets until there is at least 100 µM protein in solution with 100 mM NaCl (**Fig. 4B**). For EWS^LCD,7YS^ phase separation is not observed at the highest protein (150 µM) and salt concentrations (150 mM) tested (**Fig. 4C**). Brightfield images confirm that the increased OD_600_ is due to droplet formation. Pelleting assays performed under a single condition where EWS^LCD^ forms phase separated droplets (50 µM protein, 150 mM NaCl) reveal that single tyrosine mutations will slightly increase the amount of soluble EWS^LCD^ detected, but mutation of Y172S or the double mutant Y170S/Y172S enhance the overall solubility of EWS^LCD^ (**Fig. 4D**). Mutation of multiple tyrosines in the 7YS or 13YS promote greater than 90% soluble EWS^LCD^ under these conditions. Co-existence slab simulations highlights equilibration between dilute and dense phase for the mutants and WT at 320 K (**Fig. 4E**). The *C*_sat_ extracted from simulations show a similar trend to the pelleting assays, namely that mutating tyrosine leads to an increase in the *C*_sat_, with a greater effect observed when multiple tyrosine are mutated (**Fig. 4F)**. These experimental and simulation results suggest that tyrosine interactions with itself and other residues are important and play a crucial role in the phase separation of the EWS^LCD^.

**Figure 4.**
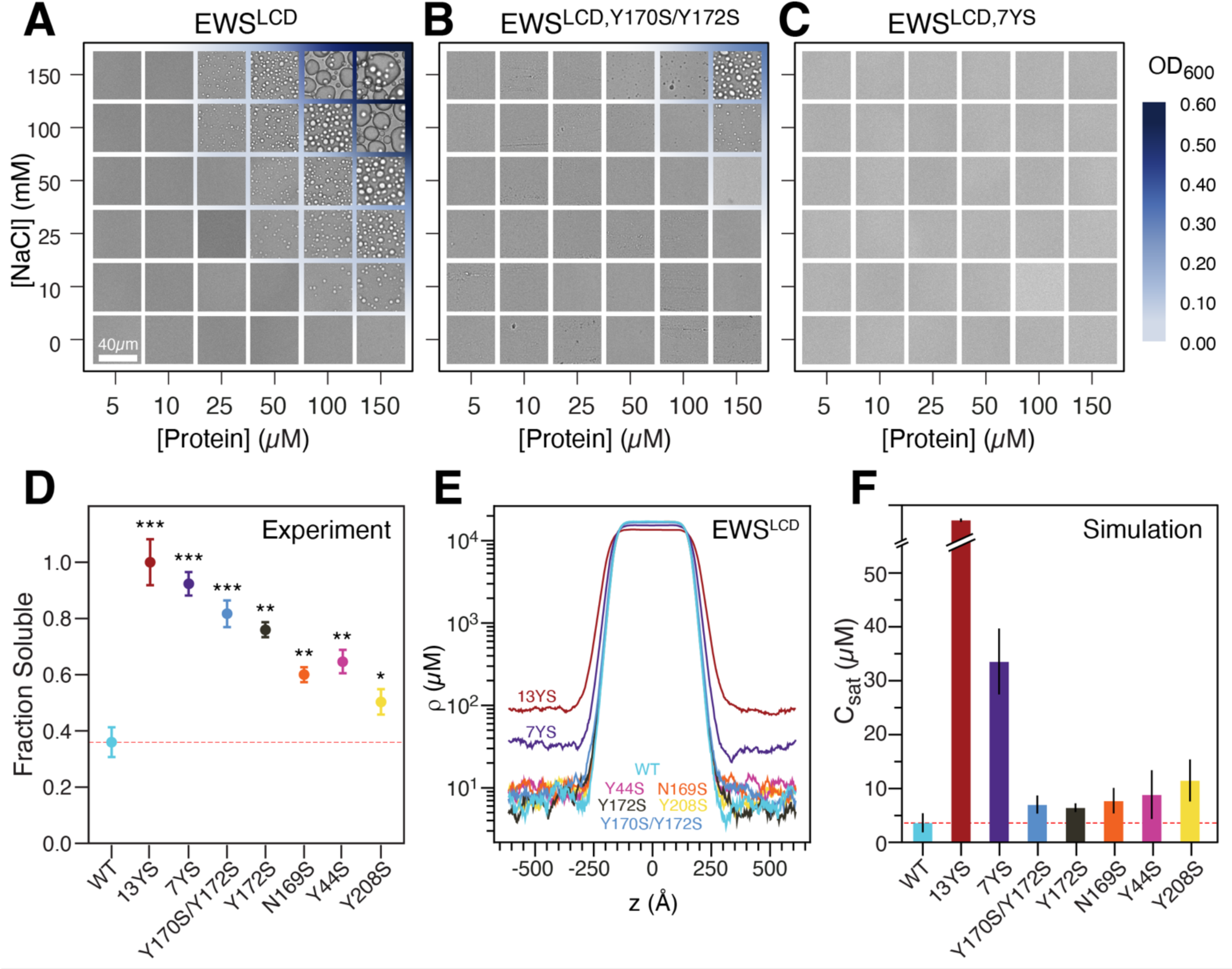
Phase separation diagrams of (**A**) EWS^LCD^, (**B**) EWS^LCD,Y170S/Y172S^, and (**C**) EWS^LCD, 7YS^, the blue gradient corresponds to increasing OD_600_. (**D**) Fraction of 50 μM EWS^LCD^ (mutants) soluble after phase separation induced with 150 mM NaCl. Significance of the change is relative to the EWS^LCD^ WT. All samples were in 20 mM Tris pH 7.4. (**E**) Density profiles along the z-dimension of the slab geometry, and (**F**) *C*_sat_ values calculated from CG coexistence simulations.

### Tyrosine Residues Affect the Dynamics of EWS^LCD^

To gain further insights into the changes in dynamics as compared to WT, we used NMR to probe the effect of tyrosine mutations on EWS^LCD^ self-association. The ^1^H,^15^N HSQCs of EWS^LCD^ and EWS^LCD,7YS^ overlay with one another well, do not indicate non-specific line broadening or degradation, enabling transfer of ∼ 75 % of the assignments from EWS^LCD^ to EWS^LCD,7YS^ spectra (**Supp. Fig. 5**). The *R*_1_, *R*_2_ rates, and hetNOEs for EWS^LCD,7YS^ are not markedly different than those measured for the EWS^LCD^ with an average *R*_1_ of ∼1.5 s^−1^, *R*_2_ of ∼3 s^−1^, and hetNOE of −0.16, but show a ∼ 1 s^−1^ decrease in *R*_2_ at Y170 compared with EWS^LCD^ (**Supp. Fig. 6A**). The relaxation rates extracted from all atom simulations show a similar qualitative trend albeit with a much larger decrease in *R*_2_ near Y170 for both EWS^LCD,Y170S/Y172S^ and EWS^LCD,7YS^ mutants (**Supp. Fig. 6B**). Intramolecular PREs from the N-terminus of the EWS^LCD,7YS,S40C^ mutant are slightly elevated when compared with the solvent PREs but overall smaller in magnitude (5-8 s^−1^) than those measured for the EWS^LCD^ (10-15 s^−1^) (**Fig. 5A**). Conversely, when the spin-label is at the C-terminus of EWS^LCD,7YS,S260C^ the magnitude of the observed PREs are similar to what was recorded for EWS^LCD,260C^ (5-10 s^−1^), at least in the region from 150-264. In the N-terminal half (1-150) of EWS^LCD,7YS,S260C^ the PREs are on average ∼2 s^−1^ slower than what was measured for EWS^LCD,260C^ (**Fig. 5A**). Overall, the magnitude of the observed PREs are prominently reduced when the spin label is within the region containing Y to S mutants (S40C vs S260C) (**Fig. 5B**). Comparison of the intramolecular pair-wise residue type contacts for EWS^LCD,Y170S/Y172S^ and EWS^LCD,7YS^ extracted from single-chain atomistic simulation shows the number of contacts decreases as more tyrosine residues are mutated to serine (**Fig. 5C**). Additionally, AUC reveals that EWS^LCD,7YS^ ensembles adopt more compact conformations when the pH is decreased from 11 to 7.4, like the behavior of EWS^LCD^ (**Fig. 5D**). Consistently, radius of gyration (*R*_g_) distribution calculated from simulations show a small expansion in EWS^LCD,7YS^ as compared to EWS^LCD^ (**Fig. 5E)**. Taken together, these findings suggest that on mutating tyrosine, intramolecular interactions within EWS^LCD^ persist; however, their frequency is reduced, resulting in a diminished capacity to promote the self-associations necessary for phase separation.

**Figure 5.**
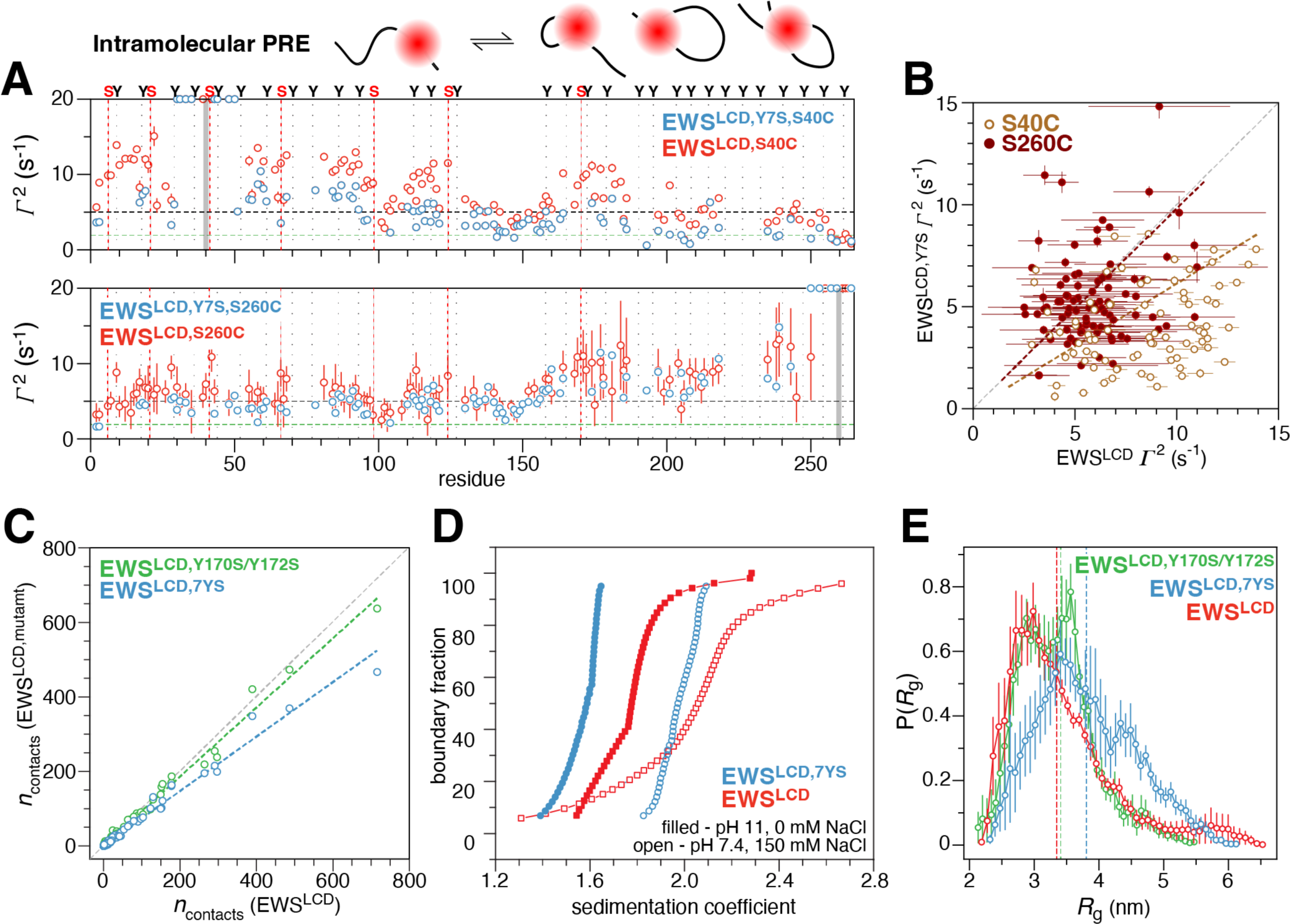
Intramolecular PRE rates (cartoon) measured for (A) tyrosine mutant EWS^LCD,7YS^ (blue) compared with PRE rates for EWS^LCD^ (red) with spin label placement (grey bar), position of tyrosine residues (dotted lines), serine substiutions (red dotted lines), and average solvent PRE (green dashed line) indicated. Black dashed line is drawn at 5 s^−1^ to guide comparison between panels. (B) Correlation of PRE rates measured for EWS^LCD,7YS^ and EWS^LCD^ with the spin label at residue 40 (brown) or 260 (maroon). (C) Correlation of average residue type pairwise contacts in EWS^LCD,7YS^ (blue) and EWS^LCD,Y170S/Y172S^ (green) extracted from single chain atomistic simulations. (D) Combined van Holde-Weischet plots of EWS^LCD^ (red) and EWS^LCD,7YS^ (blue) in phase separating (20 mM Tris pH 7.4, 150 mM NaCl, open circles) and non-phase separating conditions (20 mM CAPS pH 11, filled circles). (E) Distribution of *R*g for EWS^LCD^ (red), EWS^LCD,7YS^ (blue), and EWS^LCD,Y170S/Y172S^ (green) calculated from single chain atomistic simulations. Vertical dashed lines indicate mean *R*g.

### Condensed Phase Intermolecular Contacts Mimic Intramolecular Interactions in the Dilute Phase

Given that earlier research has suggested potential links between the condensed and dilute phases, we were motivated to delve into the interactions occurring within the condensed phase and their potential relationship with the dilute phase^64^. Investigating the condensed phase of EWS^LCD^ experimentally proved to be challenging. Given the successful utilization of condensed phase simulations in prior research^44, 62, 65^, we opted to conduct atomistic simulations of the condensed phase. Our simulations utilized a slab geometry containing 25 EWS^LCD^ chains, explicit water, and ions (details in Methods) (**Fig. 6A**). We excluded the initial 200 nanoseconds as an equilibration period for our analysis of the concentrated phase based on the *R*_g_ autocorrelation function which decays within this time for all the chains (**Supp. Fig. 7A**). Upon analyzing pairwise contact map, we observed the presence of interactions in both the N- and C-terminal regions (**Fig. 6B**). A strong correlation was noted between the pairwise residue type contacts in both the condensed and dilute phases (**Fig. 6C**). Furthermore, the pattern observed in the per-residue 1D contact propensity resembled that of the dilute phase, with peaks primarily occurring at Tyr residue positions (**Fig. 6D**). We also noticed a significant correlation for *R*_2_, indicating slower dynamics in the C- and N-terminal regions compared to the linker region (**Supp. Fig. 7B**). The *R*_2_ values in the dense phase are faster than those dilute phase which suggests slower dynamics can be attributed to the increased intermolecular interactions in condensed phase. Similar to their role in the dilute phase, we observe that several residue type pairs such as YY, QY, and QQ also contribute to stabilizing the condensed phase (**Supp. Fig. 7C**). This highlights the significance of transient interactions that induce the collapse of EWS^LCD^ in the dilute phase, also play a role in driving its phase separation.

**Figure 6.**
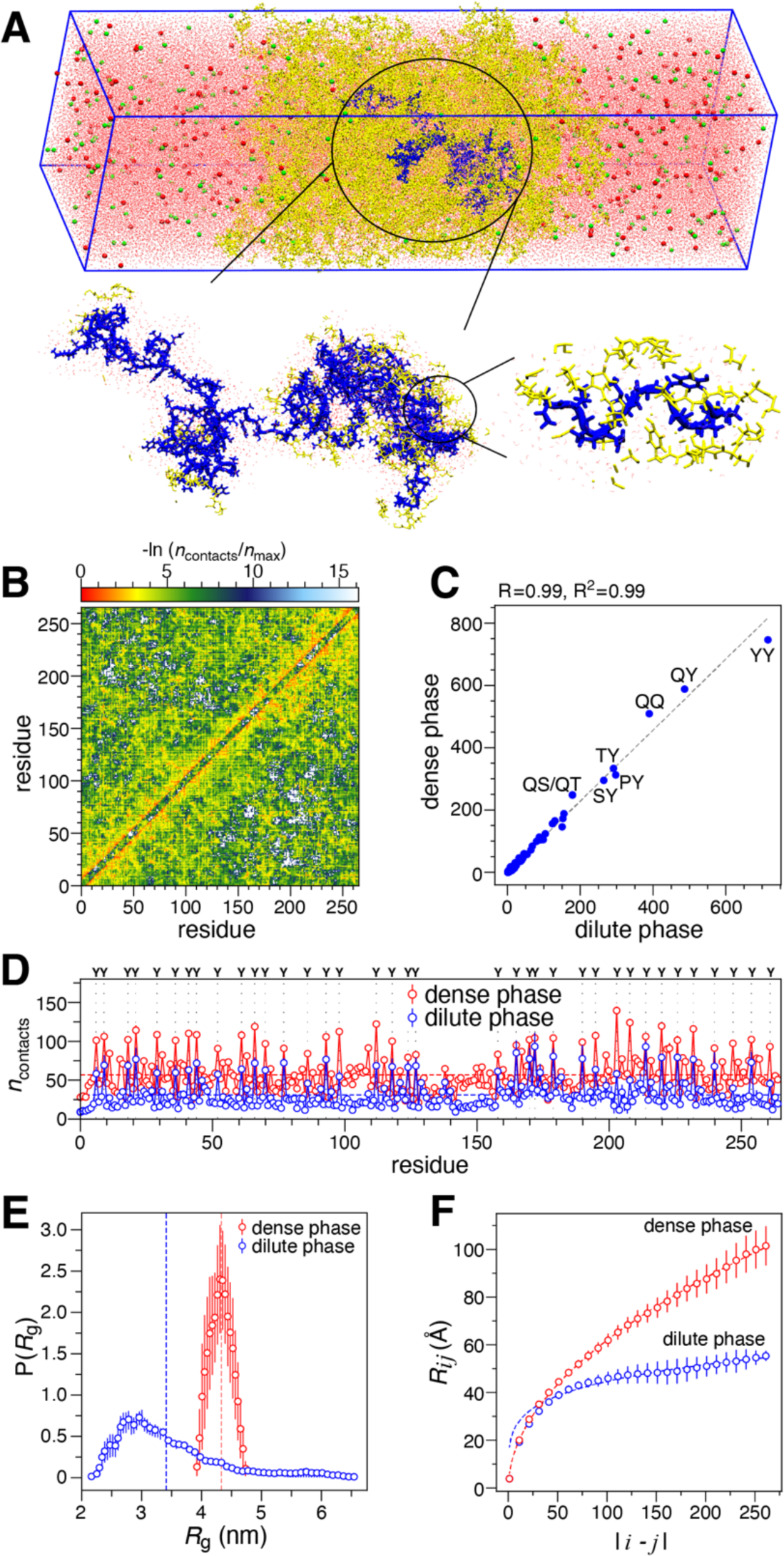
Atomistic condensed phase simulation of (**A**) 25 chains of EWS^LCD^ (yellow), with water (pink), Na (green), and Cl (red) ions. Zoomed image of one EWS^LCD^ chain (blue) and surrounding atoms. (**B**) Normalized average pairwise van der Waals contacts per EWS^LCD^ chain, and (**C**) correlation between average residue type pair contacts in the dilute and dense phase. (**D**) Average contacts from dilute (per-residue, blue) and condensed phase (per chain, red) atomistic simulations, position of tyrosine residues indicated by dotted lines. (**E**) Distribution of *R*g, and (**F**) inter-residue distances as a function of residue separation |*i-j*|, for EWS^LCD^ in dilute (blue) and dense phase (red).

We also asked how the protein conformations change when transitioning from dilute to dense phase. Notably, the *R*_g_ and scaling of average intramolecular distances (*R*_ij_) with sequence separation |i − j| shows that the chains exhibited expansion in the condensed phase compared to the dilute phase (**Fig. 6E,F**). This is consistent with expectation from homopolymer solution theories, as the protein chains inside the condensed phase can form intermolecular contacts that are same as intramolecular contacts^66^. Overall, the atomistic condensed phase simulations strongly suggest that the intramolecular interactions leading to the collapse of the C-terminal and N-terminal regions in the dilute phase also contribute to the phase separation of EWS^LCD^ via intermolecular interactions.

## Discussion

The functional basis for the differences in molecular size and sequence composition between the FET family protein LCDs is not well understood. Indeed, much of the knowledge regarding the EWS^LCD^ is surmised from studies of the FUS^LCD^, leaving a significant knowledge gap. Here, our focus was to gain insights into the molecular mechanism underlying the self-associative and phase separation properties of EWS^LCD^ given that sequence composition and amino acid patterning clearly distinguish it from both FUS^LCD^ and TAF15^LCD^. To the best of our knowledge, our study represents one of the pioneering efforts to probe the structural details of EWS^LCD^ at atomic resolution and elucidate the interactions that promote self-association and phase separation.

Through a combination of NMR, PRE analysis, and atomistic simulation, we develop an ensemble picture of the structure of EWS^LCD^. Hydrodynamic measurements using SEC and AUC indicate compaction of EWS^LCD^, and atomistic simulations reveal the C-terminal half of EWS^LCD^ preferentially populates more compact conformations when compared with the N-terminal half. Moreover, a linker region that is devoid of tyrosine residues, predominantly maintains an extended configuration. This domain-like conformational ensemble behavior is further evidenced from transverse relaxation rates which were fastest for C-terminal residues compared with N-terminal residues, and measurably slower within the linker indicating higher residue flexibility. Additionally, elevated PRE rates were observed when spin labels introduced within the N-terminal or C-terminal regions compared with the PRE rates arising from spin-labels placed in the central linker region. In contrast to previous studies on FUS^LCD^, which demonstrated transient intra- and intermolecular interactions distributed throughout the LCD without forming any compact structure^48, 67^, our observations clearly highlight the distinct behavior of EWS^LCD^. However, it is important to note that, like FUS^LCD^, EWS^LCD^ remains disordered and none of our biophysical observations indicate stabilization of secondary structure elements^67^. The ensemble-level compactness driven of the C-terminal domain observed here is consistent with previously studies that used truncation mutants or mutations of hydrophobic residues to demonstrate that the C-terminus is the most potent for inducing neoplastic transformation^8, 10^. It is worthwhile to note that the EWS^LCD^ C-terminal region is the most FUS-like in terms of sequence composition and amino acid patterning, providing a basis for commonality of function between the EWS and FUS LCDs in oncogenic fusion proteins.

Previous work determined that phase separation of FUS^LCD^ involves the formation of transient interactions involving various residue types^17, 62, 68^. Using single-chain atomistic simulations we found that the C-terminal region in EWS^LCD^ exhibits the highest intramolecular interactions followed by the N-terminal region and the linker region. This order of interaction strength can be attributed to the distinct sequence characteristics of each region. Our results show that the residue type pairs such as Tyr-Tyr, Tyr-polar (including Gln, Ser, and Thr), Tyr-Pro, and polar-polar, play pivotal roles in governing intramolecular interactions (**Fig. 7**). Mutations of EWS^LCD^ substituting aromatic tyrosines for polar serines underscore the significance of hydrophobic interactions in phase separation, with tyrosine residues playing a primary role. The biological importance of tyrosine residues in EWS^LCD^ in the context of Ewing sarcoma has been noted for some time^8, 9^. Our study highlights the importance of the specific positioning of tyrosine, such as Y170 and Y172, which when mutated to serine increased the measured *C*_sat_ resulting in a more soluble mutant of EWS^LCD^. These results are consistent with other studies that have shown that the patterning of interaction sites (no only aromatic residues) affects phase separation properties^69, 70^. Recently, Nosella and co-workers found that O-linked-N-acetylglucosaminylation reduces the propensity for EWS^LCD^ to phase separate^13^. Since 27 of the 37 tyrosines in EWS^LCD^ are immediately adjacent to an O-linked-N-acetylglucosaminylation site, the addition of a sugar moiety could reduce the probability for π-π contact formation, altering the valency and spacing of π-contacts in EWS^LCD^, and thus tuning phase separation propensity.

**Figure 7.**
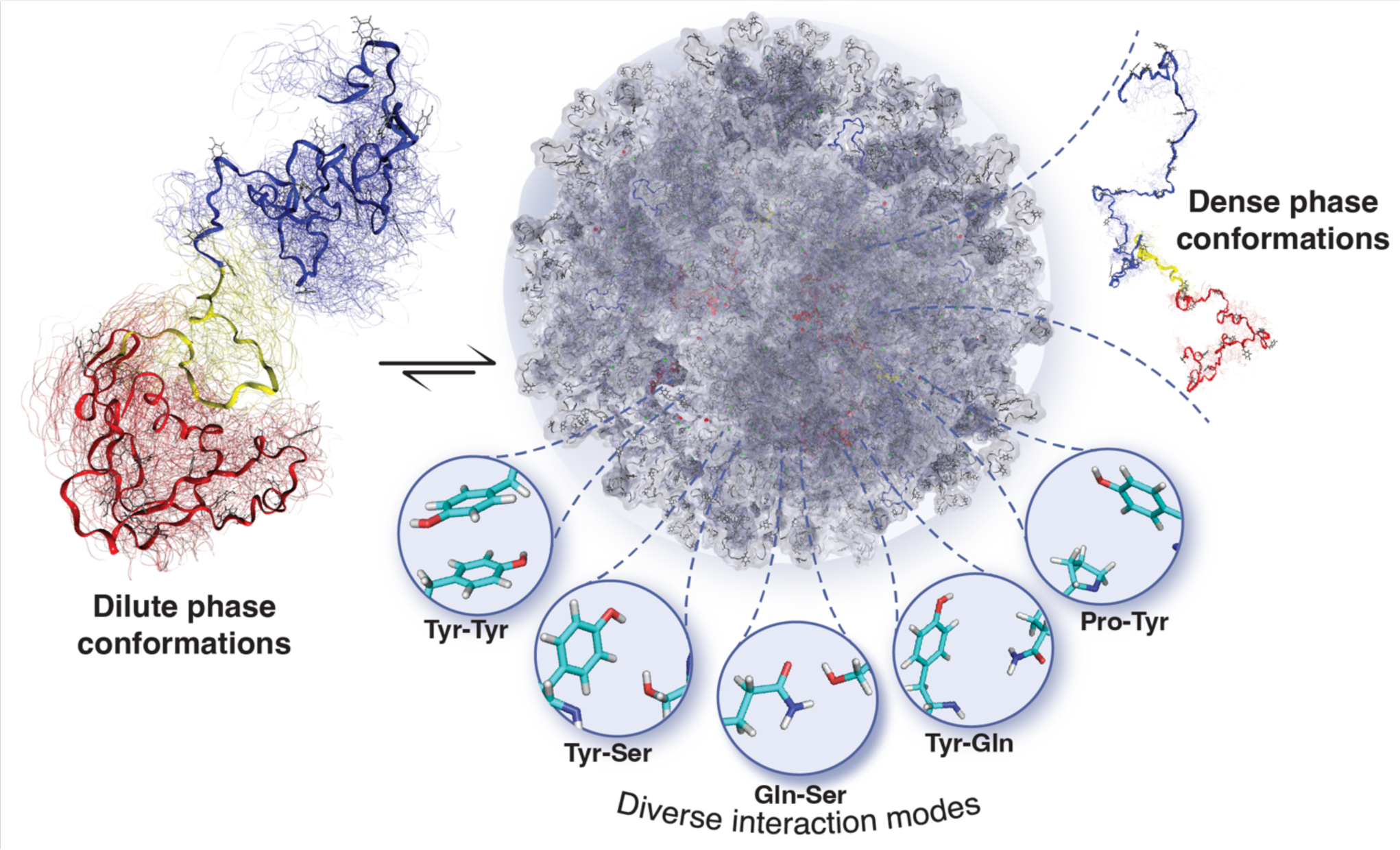
Model of EWS^LCD^ phase transition highlighting the relevant intra-residue contacts.

Atomistic simulations conducted in condensed phase demonstrate that the same residue types responsible for compaction in a dilute phase also serve to stabilize the condensed phase. These residue pairs engage in transient interactions via π-π, *sp*^2^/π, and hydrogen bonding interactions, thereby reinforcing the stability of the condensed phase^17, 62^. In addition, simulations reveal that EWS^LCD^ chains undergo expansion in the condensed phase, a phenomenon that aligns with polymer physics principles^66,71^. Single point mutations changing tyrosines and immediately adjacent residues to serines measurably altered phase separation propensity of EWS^LCD^, consistent with the behavior of single point mutations in the FUS^LCD 72^. Surprisingly, mutating the residue adjacent Y170, N169, to serine created a more soluble EWS^LCD^ mutant, supporting the hypothesis that in addition to aromatic residues, the immediate chemical environment plays an important role in the molecular grammar governing phase separation^16, 17, 62^. For example, a previous study of the phase separation of SH3_3_-FUS in the presence of PRM_4_ emphasized the significance of the quantity of tyrosine rather than its specific placement in FUS^73^. In contrast, our study reveals that both the number of tyrosine residues and their relative positioning are essential factors governing LLPS in EWS^LCD^. These results further highlight the distinct behavior of EWS^LCD^ compared to other members of the FET family. Together, our simulations and mutational analysis offer detailed molecular insight into the behavior of EWS^LCD^ within the condensed phase, and emphasize further exploration of point mutations and post-translational modifications is critical for understanding the native and oncogenic functions of EWS and EWS::FLI1 respectively.

Recently, Chong et al. proposed a yin-yang hypothesis that optimal interactions involving LCD-LCD contacts within the EWS::FLI1 fusion are responsible for activating the transcription of the oncogene, yet phase separation of EWS::FLI1 leads to gene repression^18^. Further, the transformative properties of EWS::FLI1, including binding GGAA-microsatellites and subsequent recruitment of the BAF complex and other chromatin modifiers have been suggested to rely on phase separation^9, 15^. While the self-associative, and possibly phase separation properties of EWS::FLI1 are clearly important in oncogenesis, the multivalent nature of the EWS^LCD^ is also important for native EWS functions such as mediating protein-protein and protein-RNA interactions important for centromere maintenance^74^. Together, these studies demonstrate that EWS^LCD^-mediated self-association is central to the native functions of EWS and aberrant functions of EWS::FLI1, however, molecular and structural insights were missing. The results presented here provide the necessary insight into the structure and function of EWS^LCD^ in both dilute and condensed phases and begin to unravel the molecular basis for EWS and EWS::FLI1 biological functions. Our characterization of EWS^LCD^ self-association and phase separation contrasts with observations of the behavior of FUS^LCD^ and provides new insights and further stratifies the functional differences of the FET family proteins. Finally, this work supports the importance of understanding the heterotypic interactions and emergent functions of biomolecular condensates and forms a foundation for future studies aimed at understanding how modulating phase separation alters cellular activities.

## Abbreviations

EDTA: ethylenediaminetetraacetic acid
EWS: RNA-binding protein EWS
EwS: Ewing sarcoma
FRAP: fluorescence recovery after photobleaching
FUS: fused in sarcoma
HSQC: heteronuclear single quantum coherence
IDR: intrinsically disordered region
LCD: low-complexity domain
LLPS: liquid-liquid phase separation
NMR: nuclear magnetic resonance
PRE: paramagnetic relaxation enhancement
SEC: size-exclusion chromatography
TATA-binding protein-associated factor 2N TAF15

## Acknowledgements

DSL is the Shohet Family Fund for Ewing Sarcoma Research St. Baldrick’s Scholar and acknowledges the support of the St. Baldrick’s Foundation (634706). This study was funded in part by the Welch Foundation AQ-2001-20190330 (to DSL) and A-2113-20220331 (to JM), NIGMS R01GM140127 (to DSL) and R01GM136917 (to JM), and GCCRI Startup Funds (DSL). The authors thank Dr. Dmitri Ivanov for generous access to an imagining microwell plate reader instrument. The TOC graphic was created with BioRender.com. We gratefully acknowledge the computational resources provided by the Texas A&M High Performance Research Computing (HPRC). This work is based upon research conducted in the Structural Biology Core Facilities, a part of the Institutional Research Cores at the University of Texas Health Science Center at San Antonio supported by the Office of the Vice President for Research and the Mays Cancer Center Drug Discovery and Structural Biology Shared Resource (NIH P30 CA05417X4).

## Associated Content

Two supplemental table describing the construction and purpose of the mutants used in this study as well as seven additional figures are available in the Supplemental Materials file.

## Data Availability

NMRPipe processing scripts are available upon reasonable request, expression plasmids containing the EWS^LCD^ wildtype and mutants were deposited with Addgene (accession numbers to be added).

## Author Contributions

CNJ collected data, analyzed data, wrote manuscript, KS collected data, analyzed data, wrote manuscript, ES collected data, analyzed data, AKMR collected data, AB collected data, XX collected data, JM analyzed data, wrote manuscript, obtained funding, supervised research, DSL analyzed data, wrote manuscript, obtained funding, supervised research.

