## Supplemental Materials for "Insights into Molecular Diversity within the FET Family: Unraveling Phase Separation of the N-Terminal Low Complexity Domain from RNA-Binding Protein EWS"

**Supplemental Table 1:** Primers and template synthetic codon-optimized genes used to construct EWS$^{LCD}$ mutants.

| **Construct** | **Template** | **Forward Primer** | **Reverse Primer** |
|---|---|---|---|
| EWS$^{LCD,S40C}$ | EWS$^{LCD}$ | GCCAGCAGTGCTATGGTACCTA TGGCCAGCCGACCG | CCATAGCACTGCTGGCCATACG CCTGGGTGGTCTGGG |
| EWS$^{LCD,S116C}$ | EWS$^{LCD}$ | GCCCAGTGCGCCTATGGTACCC AGCCGGCATATCC | TAGGCGCACTGGGCCGCATAG CTTGCCTGGGTGG |
| EWS$^{LCD,S149C}$ | EWS$^{LCD}$ | CGAAACCTGCCAGCCGCAGAGC AGCACCGGCG | GGCTGGCAGGTTTCGGTCGGTT TGTTACCATCCTGCGG |
| EWS$^{LCD,S239C}$ | EWS$^{LCD}$ | GCCGACCTGCTATCCGCCGCAG ACCGGTAGCTATAGC | GGATAGCAGGTCGGCGGCTGC TGACCATAGCTGCTTTG |
| EWS$^{LCD,S260C}$ | EWS$^{LCD}$ | AGCTCTTGTTATGGCCAGCAGTG AGGATCCGGCTGC | GGCCATAACAAGAGCTCTGCT GGCTATACTGGCTCGG |
| EWS$^{LCD,Y44S}$ | EWS$^{LCD}$ | GGTACCTCTGGCCAGCCGACCG ATGTGAGCTATACCCAGGC | GCTGGCCAGAGGTACCATAGC TCTGCTGGCCATACGCCTGGG |
| EWS$^{LCD,Y170S}$ | EWS$^{LCD}$ | AGCAATTCTAGCTATCCGCAGG TTCCGGGCAGCTATCCG | GATAGCTAGAATTGCTCTGGCC ATAACCCAGGCTCGGCTG |
| EWS$^{LCD,Y172S}$ | EWS$^{LCD}$ | TTATAGCTCTCCGCAGGTTCCGG GCAGCTATCCGATGCAGCC | CTGCGGAGAGCTATAATTGCTC TGGCCATAACCCAGGCTCGG |
| EWS$^{LCD,Y208S}$ | EWS$^{LCD}$ | AGCAGCTCTAGCCAGCAGAACA CCTATGGTCAGCCGAGCAGCTA TGGCC | GCTGGCTAGAGCTGCTCTGATC ATAGCTGGTCGGCTGGGTGCTG C |
| EWS$^{LCD,Y170S/Y172S}$ | EWS$^{LCD,Y170S}$ | TTCTAGCTCTCCGCAGGTTCCGG GCAGCTATCCGATGCAGCC | CTGCGGAGAGCTAGAATTGCT CTGGCCATAACCCAGGCTCGG |
| EWS$^{LCD,N169S}$ | EWS$^{LCD}$ | CAGAGCAGCTATAGCTATCCGC AGGTTCCGGGC | GCTATAGCTGCTCTGGCCATAA CCCAGGCTCGGC |

**Supplemental Table 2:** Mutants used to investigate EWS$^{LCD}$ phase separation.

| Construct | Mutation(s) | Purpose |
|---|---|---|
| EWS$^{LCD, S40C}$ | S40C | For attachment of PRE label in tyrosine intact EWS$^{LCD}$ |
| EWS$^{LCD, 2116C}$ | S116C | For attachment of PRE label in tyrosine intact EWS$^{LCD}$ |
| EWS$^{LCD, S149C}$ | S149C | For attachment of PRE label in tyrosine intact EWS$^{LCD}$ |
| EWS$^{LCD,S239C}$ | S239C | For attachment of PRE label in tyrosine intact EWS$^{LCD}$ |
| EWS$^{LCD,S260C}$ | S260C | For attachment of PRE label in tyrosine intact EWS$^{LCD}$ |
| EWS$^{LCD, Y170S/Y172S}$ | Y170S, Y172S | To investigate the effect on phase separation by mutating tyrosines in close proximity |
| EWS$^{LCD, 7YS}$ | Y6S, Y21S, Y41S, Y66S, Y98S, Y124S, Y170S | To investigate the effect on phase separation mutating multiple tyrosine residues in the N-terminus |
| EWS$^{LCD,13YS}$ | Y179S, Y190S, Y195S, Y203S, Y208S, Y214S, Y220S, Y226S, Y232S, Y240S, Y247S, Y254S, Y261S | To investigate the effect on phase separation from mutating multiple tyrosine residues in the C-terminus |
| EWS$^{LCD, Y172S}$ | Y172S | To investigate the effect on phase separation from mutating a single tyrosine in close proximity to another tyrosine |
| EWS$^{LCD,N169S}$ | N169S | To investigate the effect on phase separation from mutating a non-tyrosine residue |
| EWS$^{LCD,Y44S}$ | Y44S | To investigate the effect on phase separation from mutating a single tyrosine in the N-terminus |
| EWS$^{LCD, Y208S}$ | Y208S | To investigate the effect on phase separation from mutating a single tyrosine in the C-terminus |
| EWS$^{LCD,7YS, S40C}$ | Y6S, Y21S, S40C, Y41S, Y66S, Y98S, Y124S, Y170S | To investigate how mutating multiple tyrosine residues in the N-terminus (non-SYGQ repeat) affects inter and intra residue interactions in PRE experiments. S40C was used for attachment of PRE label. |
| EWS$^{LCD,7YS, S260C}$ | Y6S, Y21S, Y41S, Y66S, Y98S, Y124S, Y170S, S260C | To investigate how mutating multiple tyrosine residues in the N-terminus (non-SYGQ repeat) affects inter and intra residue interactions in PRE experiments. S260C was used for attachment of PRE label. |

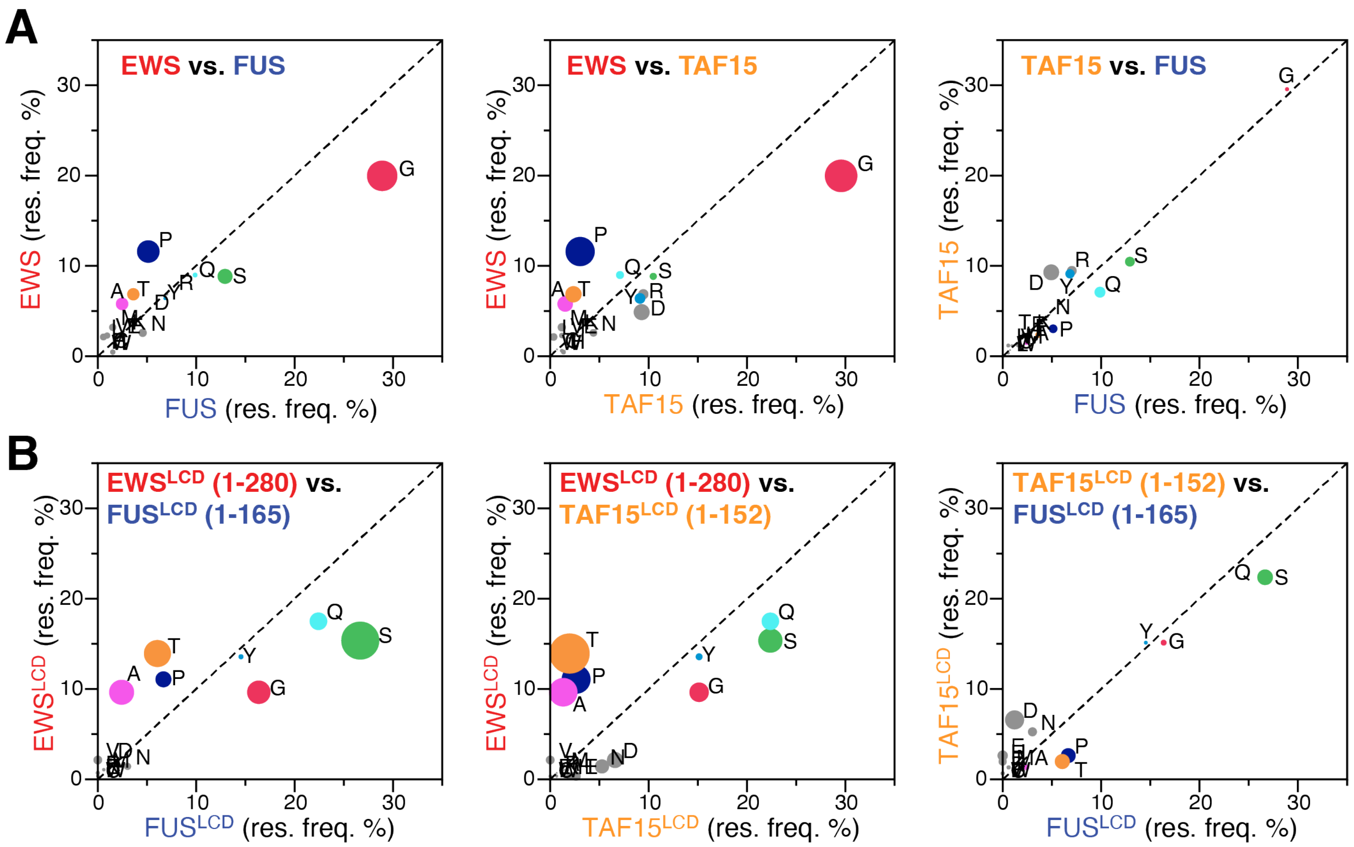


**Supplementary Figure 1:** Amino acid distribution in FET family proteins, percent abundance per amino acid where the size of the circle correlates positively with the enrichment of each residue (e.g., large red circle indicates more glycine in FUS than EWS). (**A**) Full-length EWS vs. full-length FUS (left), full-length EWS vs. full-length TAF15 (middle), and full-length TAF15 vs. full-length FUS (left). (**B**) The low complexity domains (LCD) from EWS ($EWS^{LCD}$, residues 1-280) vs. FUS ($FUS^{LCD}$, residues 1-165) (left), $EWS^{LCD}$ vs. TAF15 ($TAF15^{LCD}$, residues 1-152) (middle), and $TAF15^{LCD}$ vs. $FUS^{LCD}$ (right).

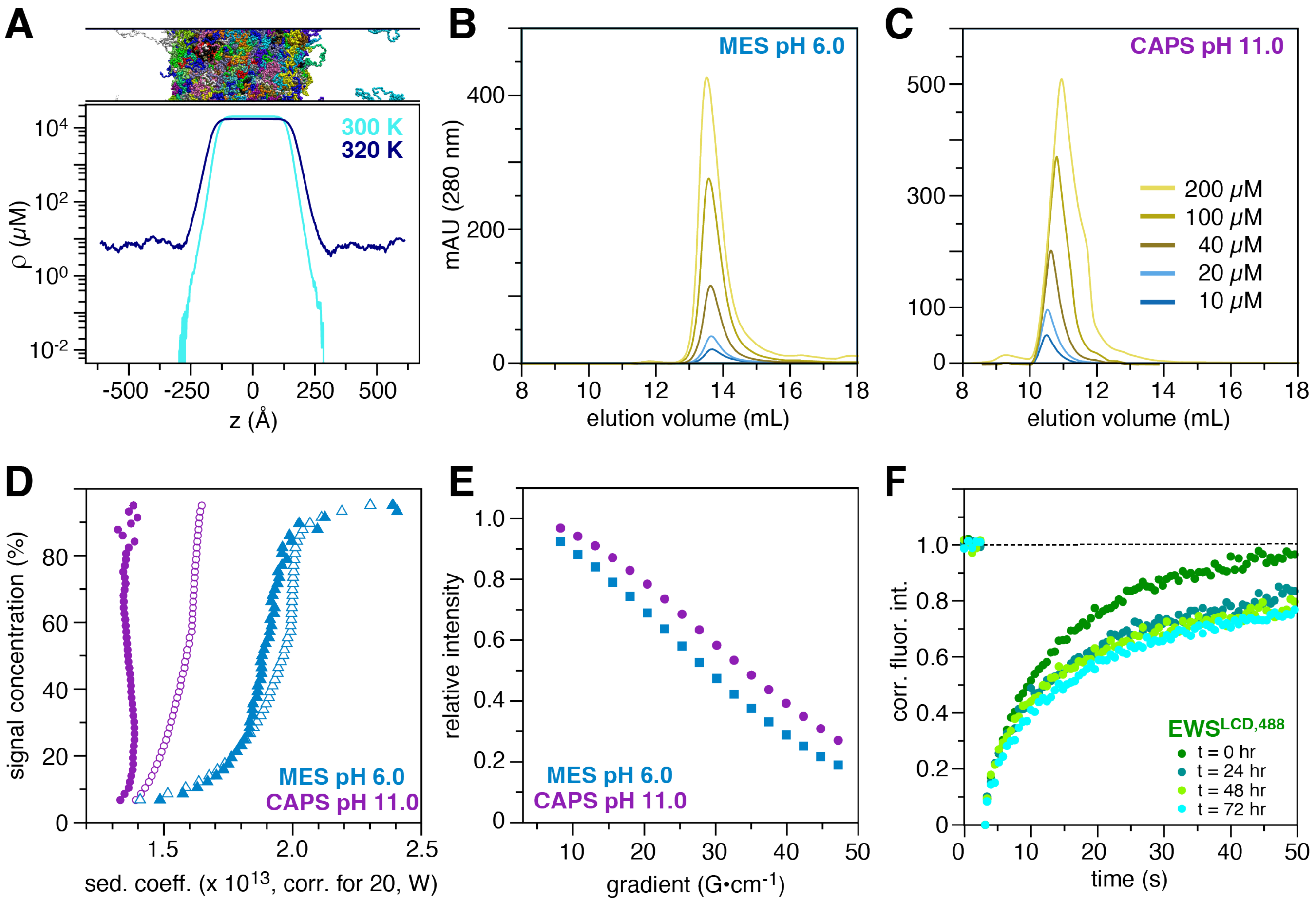


**Supplementary Figure 2:** (**A**) Concentration profile of 100 chains of $EWS^{LCD}$ from co-existence CG simulations at two temperatures. Cartoon representation of co-existence phase shown at the top. Size exclusion chromatography curves of increasing concentrations of $EWS^{LCD}$ in, (**B**) 20 mM MES pH 6.0, or (**C**) 20 mM CAPS pH 11. (**D**) Integral distributions from enhanced van Holde-Weischet plots of 25 µM $EWS^{LCD}$ (open shapes) or $EWS^{LCD,7YS}$ (filled shapes) reveal single component systems from sedimentation velocity experiments in 20 mM MES pH 6.0 or 20 mM CAPS pH 11. (**E**) Translational diffusion of $EWS^{LCD}$ measured by diffusion ordered spectroscopy in MES pH 6.0 (squares) and CAPS pH 11.0 (circles) (**G**) Time-course fluorescence recovery after photobleaching recovery curves for $EWS^{LCD,488}$.

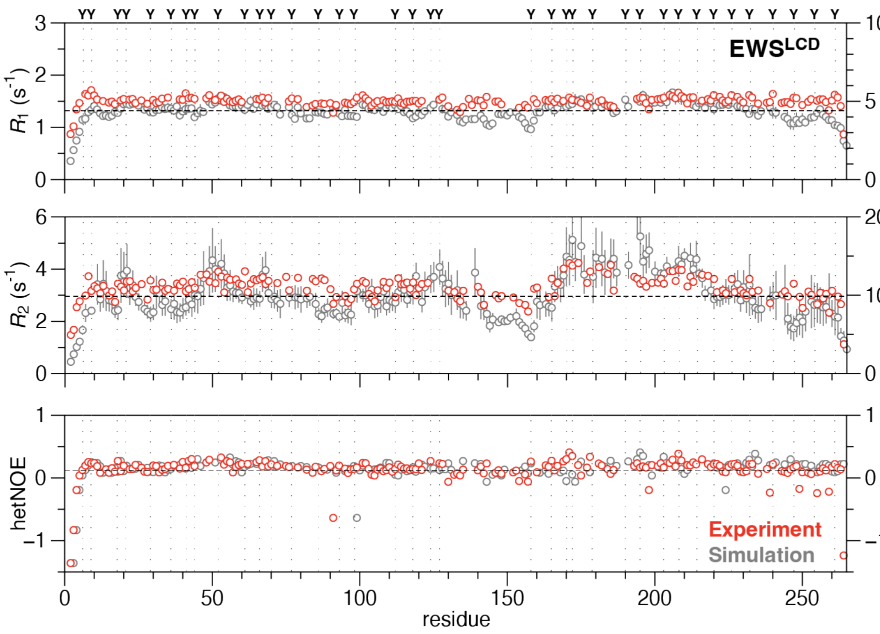


**Supplementary Figure 3:** The $^{15}$N longitudinal ($R_1$, top), $^{15}$N transverse ($R_2$, middle) relaxation rates, and $^{15}$N heteronuclear NOE values (bottom) measured on 50 µM $^{15}$N EWS$^{LCD}$ in 20 mM MES pH 5.5. The distribution of tyrosine residues is depicted above the top panel and with dotted line that extends across all panels. Experimentally measured rates (red) are compared with those computed from fitting the N-H bond autocorrelation function averaged over 100 ns blocks from single chain atomistic simulations (grey). Black dashed lines approximate average experimental (computed) values of 1.50 ± 0.01 (4.39 ± 0.21) s$^{-1}$, 3.28 ± 0.02 (9.92 ± 1.7) s$^{-1}$, and 0.13 (0.12) for $R_1$, $R_2$, and hetNOE respectively.

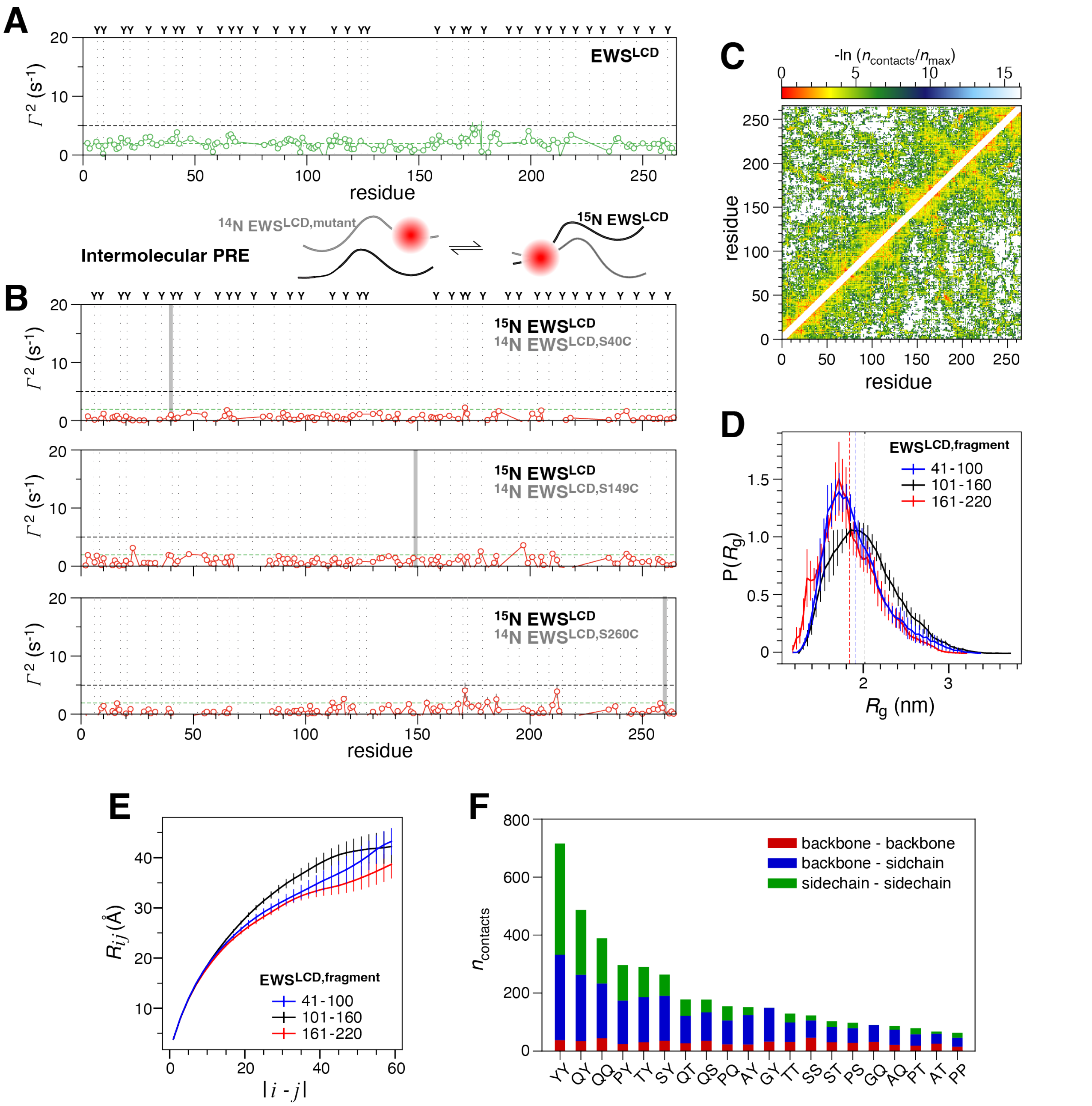


**Supplementary Figure 4:** (**A**) Solvent paramagnetic relaxation enhancement (PRE) rates plotted per residue for $EWS^{LCD}$ measured using 0.5 μM 5-MSL spin label in solution with 50 μM $EWS^{LCD}$ which contains no native cysteines. Intermolecular PREs rates (cartoon) for (**B**) $^{15}$N $EWS^{LCD}$ mixed with 10 % (w/w) spin-labled $^{14}$N $EWS^{LCD,mutant}$. Position of attached nitroxide spin label (40, 149, 260) is indicated by a gray bar. Black dotted line is an arbitrary line drawn at 5 $s^{-1}$ to help guide the eye when evaluating the plotted rates, green line indicates average solvent PRE. (**C**) Average pairwise intrachain interactions from single chain atomistic simulations. The contacts are normalized to the highest average contact. Comparison of (**D**) the *R*g distribution (vertical lines represent the mean values of the distribution), and (**E**) inter residue distance plotted as a function of residue separation |i-j| calculated from single chain atomistic simulations of three equal-length segments from $EWS^{LCD}$. (**F**) Average pairwise residue type interactions from single chain atomistic simulations separated into backbone-backbone (red), backbone-sidechain (blue), and sidechain-sidechain (green) contacts.

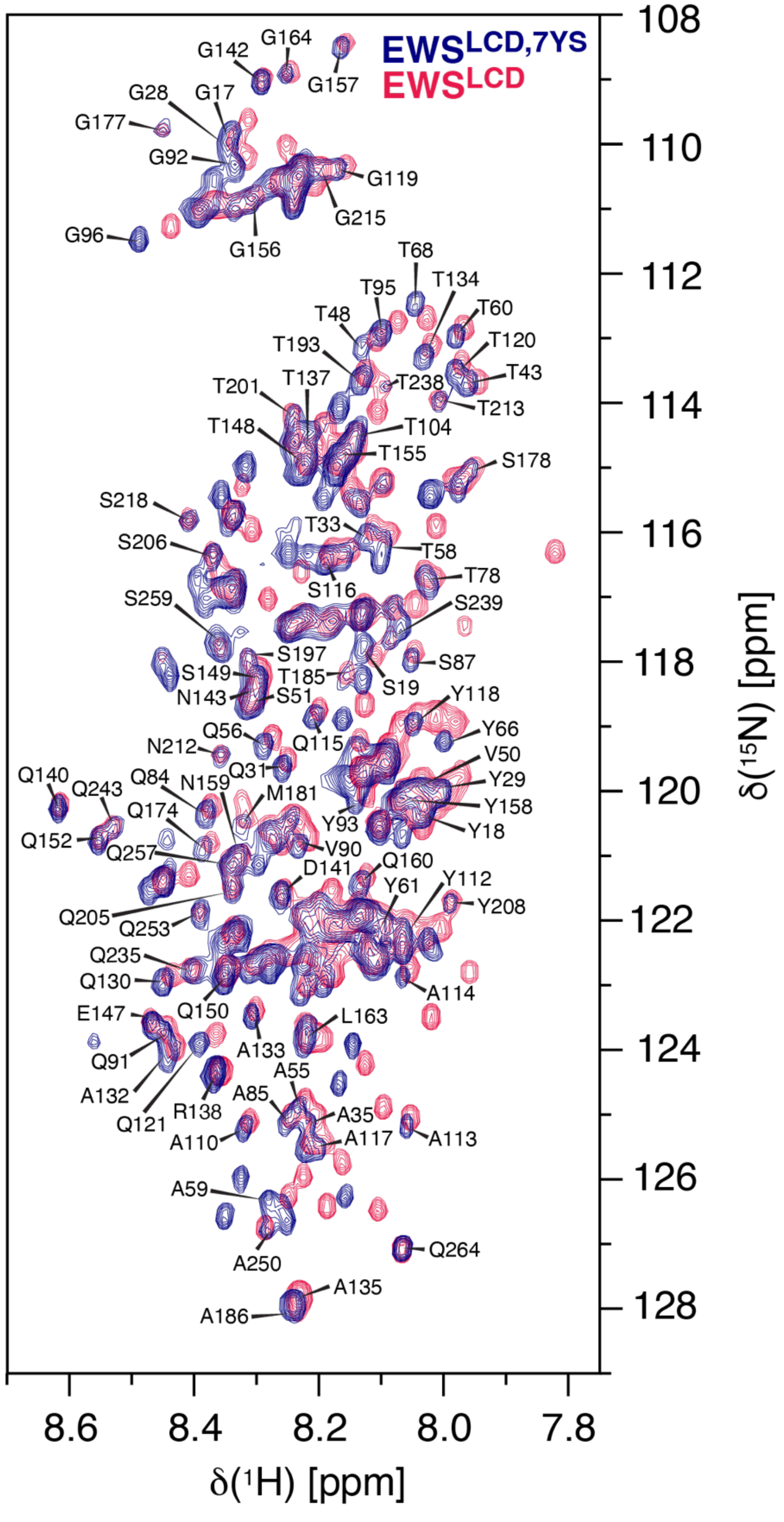


**Supplementary Figure 5:** $^{1}H,^{15}N$-HSQC spectra of 100 μM $^{13}C,^{15}N$-EWS$^{LCD}$ (red) or $^{13}C,^{15}N$-EWS$^{LCD,7YS}$ (blue) acquired at 298 K on in 20 mM MES pH 5.5, 0.5 mM PMSF, 0.2 mM EDTA, 5 % $D_2O$. Label resonances indicate assignments that were transferred from the EWS$^{LCD}$ spectrum to the EWS$^{LCD,7YS}$ spectrum.

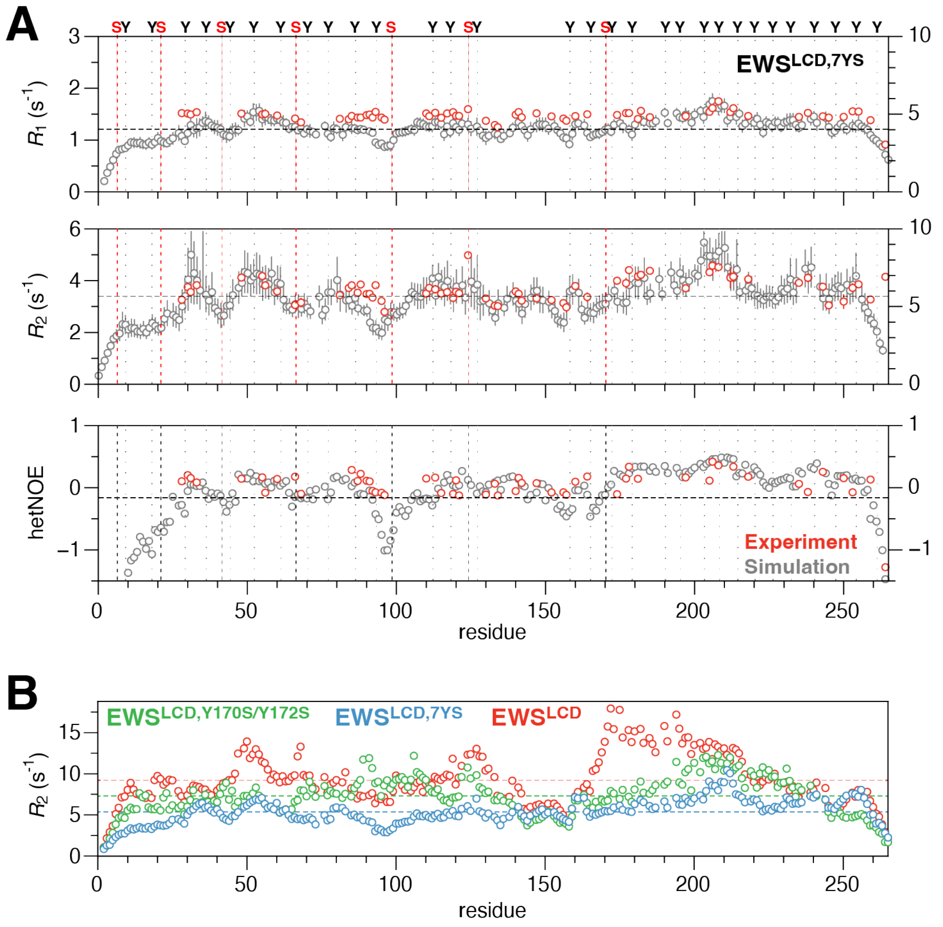


**Supplementary Figure 6:** (**A**) The $^{15}$N longitudinal ($R_1$, top), $^{15}$N transverse ($R_2$, middle) relaxation rates, and $^{15}$N heteronuclear NOE values (bottom) measured on a sample of 50 µM of $^{15}$N,$^{13}$C EWS$^{\text{LCD,7YS}}$ in 20 mM MES pH 5.5. The distribution of tyrosine residues is depicted above the top panel and with dotted line that extends across all panels. Experimentally measured rates (red) are compared with those computed from fitting the N-H bond autocorrelation function averaged over 100 ns blocks from single chain atomistic simulations (grey). Black dashed lines approximate average experimental (computed) values of 1.47 ± 0.02 (4.03 ± 0.37) s$^{-1}$, 3.68 ± 0.06 (5.66 ± 0.86) s$^{-1}$, and 0.06 (-0.16) for $R_1$, $R_2$, and hetNOE respectively. (**B**) $^{15}$N transverse relaxation rate comparison of EWS$^{\text{LCD}}$ (red), EWS$^{\text{LCD,7YS}}$ (blue) and EWS$^{\text{LCD,Y170S, Y172S}}$ (green) from single chain atomistic simulations. The rates are computed from fitting the N-H bond autocorrelation function averaged over 100ns blocks. The horizontal lines represent mean transverse relaxation values of the three systems.

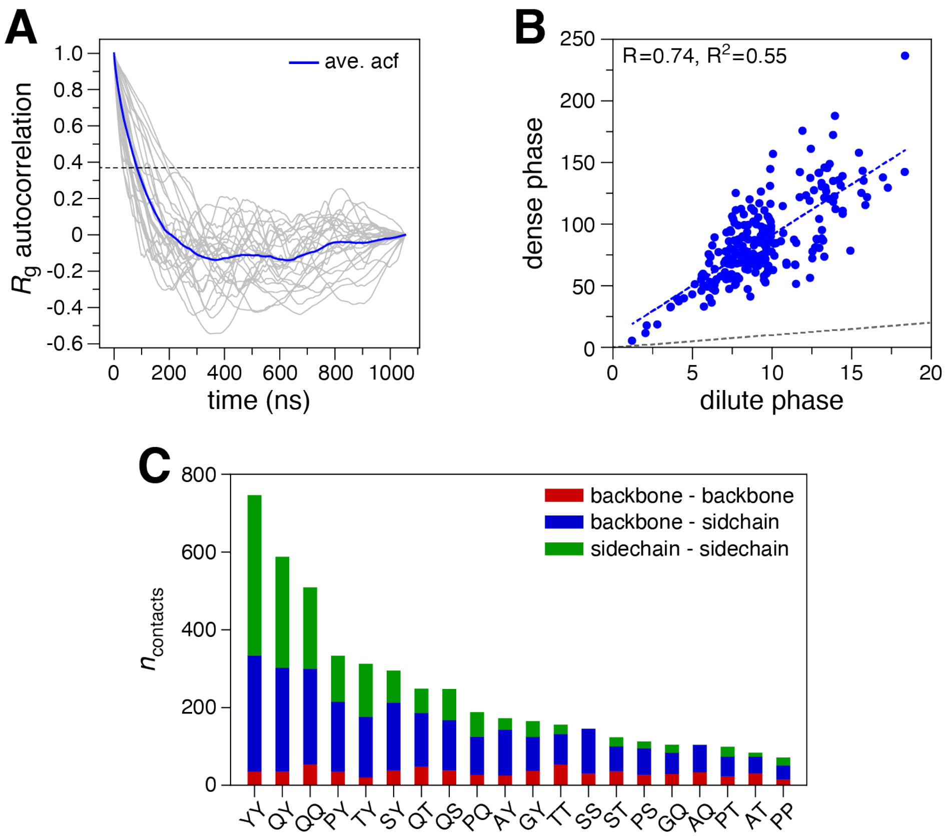


**Supplementary Figure 7:** (**A**) $R_g$ autocorrelation (acf) of 25 chains (gray) and the average (blue) reveals the correlation decays within 200 ns for all chains and therefore the first 200 ns may be excluded as equilibration for the analysis of condensed phase. (**B**) Correlation of calculated transverse relaxation rate ($R_2$) for $EWS^{LCD}$ in dilute and condensed phases. The gray dashed line represents similar transverse relaxation value in both the phases. The regression fit (blue) and the Pearson correlation coefficient (R) and its square ($R^2$) are shown at the top of the plot. (**C**) Average pairwise residue type interactions from condensed phase atomistic simulations separated into backbone-backbone (red), backbone-sidechain (blue), and sidechain-sidechain (green) contacts.
